# HIV-1 Tat favors the multiplication of *Mycobacterium tuberculosis* and toxoplasma by inhibiting clathrin-mediated endocytosis and autophagy

**DOI:** 10.1101/2025.05.02.651870

**Authors:** Aurélie Rivault, Audrey Bernut, Myriam Ben-Neji, Magali Abrantes, Maxime Jansen, Sylvaine Huc-Brandt, Sébastien Besteiro, Yann Bordat, Nelly Audemard, Margaux Mesleard-Roux, David Perrais, Olivier Neyrolles, Geanncarlo Lugo-Villarino, Christel Vérollet, Lucile Espert, Bruno Beaumelle

## Abstract

HIV-1 and *Mycobacterium tuberculosis* (Mtb) coinfections are a major public health problem but are not well characterized. HIV-1 Tat is secreted by infected cells, generating nanomolar concentrations of Tat in the sera of people living with HIV. Circulating Tat enters cells, binds to PI(4,5)P_2_ then undergoes palmitoylation, thereby becoming resident on this phosphoinositide. Here, we found that Tat favors the multiplication of Mtb in macrophages. Moreover, Tat renders zebrafish larvae more sensitive to mycobacterial infection. We found that Tat binding to PI(4,5)P_2_ and palmitoylation enable Tat to inhibit the recruitment of the AP-2 adaptor, thereby inhibiting clathrin-mediated endocytosis and in turn autophagy. This inhibition prevents the degradation of intracellular pathogens such as Mtb and opsonized *Toxoplasma gondii,* but also of lipid droplets, thereby facilitating the access of these pathogens to lipids. We thus identified a mechanism enabling HIV Tat to favor the multiplication of intracellular pathogens such as Mtb.

## Introduction

The Type 1 Human Immunodeficiency Virus (HIV-1) / *Mycobacterium tuberculosis* (Mtb) coinfection poses a major public health challenge. In 2022, approximately 650,000 HIV-1-associated deaths were reported, of which 30% involved Mtb. Tuberculosis remains the leading cause of death among people living with HIV (PLWH) [1]. The risk of developing tuberculosis is 16-to 27-fold higher for PLWH than for HIV-uninfected individuals [2–4].

The mechanisms by which HIV-1 facilitates infection by Mtb remain poorly understood [2, 5, 6]. While it is clear that CD4^+^ T-cell depletion caused by HIV-1 strongly impairs the immune response against Mtb, PLWH become more susceptible to Mtb infection shortly after seroconversion, long before peripheral CD4+ T-cell counts drop [2, 6]. Conversely, although CD4+ and CD8+ T-cell levels and functionalities are restored after approximately 12 months of antiretroviral therapy (ART) [7], PLWH under ART still remain 4-to 7-fold more susceptible to Mtb infection than the HIV-naive population [2, 4]. Moreover, *ex vivo* studies using macrophages only demonstrated a facilitating effect of HIV-1 infection on Mtb multiplication [8]. This suggests that HIV-1 enhances Mtb multiplication inside macrophages independently of other cell types, although the precise underlying mechanisms remain to be elucidated.

Autophagy is a key immune mechanism enabling cells to fight pathogens [9]. Typically, autophagy induction in cells infected by intracellular pathogens such as Mtb leads to their degradation [10] and, conversely, Mtb elimination by several antibiotics relies on autophagy [11]. The building of autophagic vesicles requires several autophagy-related (ATG) proteins and relies on several membrane sources, among which ATG16L1^+^ intermediates that result from clathrin-mediated endocytosis (CME). Accordingly, inhibiting CME blocks autophagy [12–14]. Autophagy can also degrade lipid droplets (LDs), thereby regulating their intracellular concentration [15]. LDs, which contact vacuoles containing Mtb or the parasite *Toxoplasma gondii* provide a source of lipids to these intracellular pathogens [16, 17]. Several other intracellular pathogens including bacteria, viruses and parasites induce LD formation or stabilization and exploit these structures as lipid sources to fuel their replication [18, 19].

ART is a combination of inhibitors that most often target HIV-1 enzymes, *i.e.* reverse transcriptase, integrase and protease [20]. However, the production of regulatory HIV-1 proteins such as Nef and Tat is largely unaffected by ART [21–23]. These HIV regulatory proteins may contribute to the increased susceptibility to TB observed in PLWH under ART compared to HIV-uninfected individuals.

We previously showed that HIV-Tat is strongly secreted by infected CD4^+^ T-cells, with approximately two thirds of newly synthesized Tat being secreted. This unconventional secretion occurs through the plasma membrane following Tat recruitment by PI(4,5)P_2_ [24]. Nanomolar levels of Tat are accordingly detected in the sera [23] and cerebrospinal fluid (CSF) [22, 25] of PLWH, even under ART and with viral RNA levels below 50 copies/ml, *i.e.* virally suppressed. Circulating Tat enters uninfected cells through CME, and subsequently translocates from endosomes to the cytosol [26, 27]. Once in the cytosol, Tat binds to PI(4,5)P_2_ at the plasma membrane where, in uninfected cells only, it undergoes palmitoylation [28]. This lipid anchor enables Tat to become resident on PI(4,5)P_2_ and to interfere with the recruitment by this phosphoinositide of key effectors involved in phagocytosis and neurosecretion machineries, *i.e.* Cdc42 and annexin2, respectively [28–30].

Here we show that Tat interferes with the PI(4,5)P_2_-mediated recruitment of the key CME adaptor AP-2. This results in CME inhibition that, in turn, strongly reduces the autophagic response to intracellular pathogens such as Mtb and opsonized *T. gondii*. Additionally, this autophagy inhibition protects LDs from degradation. Tat thereby promotes the multiplication of Mtb and Toxoplasma in primary human macrophages and facilitate mycobacterial replication in zebrafish larvae. By inhibiting autophagy, Tat likely supports the proliferation of intracellular pathogens both by preventing their degradation and by facilitating their access to cellular lipid stores.

## Results

### HIV-1 Tat favors the multiplication of intracellular pathogens in macrophages

We observed that HIV-1 Tat, at physiological concentration (15 nM) [23] facilitates the growth of dsRed-expressing Mtb in human monocyte-derived macrophages (hMDMs) after 3 days of infection (Figure 1A-C). While Tat moderately increased the percentage of infected cells (Figure 1A), Tat effect on mycobacterial intracellular multiplication was more pronounced (Figure 1B-C). Throughout this study we used two Tat point mutants as negative controls: Tat-W11Y, which does not enter cells efficiently [27], weakly binds PI(4,5)P_2_ [24] and is not palmitoylated, and Tat-C31S, which enters cells normally and binds to PI(4,5)P_2_, but is not palmitoylated, thus does not remain on this phosphoinositide and is instead rapidly secreted. Neither Tat-W11Y nor Tat-C31S affect PI(4,5)P_2_-dependent traffic such as phagocytosis or neurosecretion [28]. These control mutants did not significantly affect Mtb intracellular multiplication (Figure 1B-C). We wondered whether Tat effect on Mtb multiplication could be indirect, *i.e.* mediated by Tat-induced inflammatory cytokines. Tat has been shown to trigger the secretion of IL-6, IL-1β, TNF-α and IL-10 by cells from the monocyte/macrophage lineage [31–33]. These cytokines are generally considered detrimental for Mtb intracellular multiplication [34]. Although the results were donor-dependent, Tat-C31S induce cytokines by primary human macrophages as efficiently as WT Tat (Figure S1). Nevertheless Tat-C31S did not activate NF-κB (Figure S2), indicating that its capacity to trigger cytokine release could rely on the activation of other transcription factors such as AP-1 [35]. Tat-W11Y was as efficient as WT Tat in activating NF-κB (Figure S2) and even more efficient in triggering cytokine production (Figure S1). The absence of any significant effect of Tat-W11Y and Tat-C31S on Mtb multiplication (Figure 1B-C) indicates that Tat facilitating effect on Mtb intramacrophagic multiplication is unlikely mediated by cytokine release. If cytokines were responsible, Tat-W11Y would have been more efficient than WT Tat in promoting Mtb multiplication.

**Figure 1.**
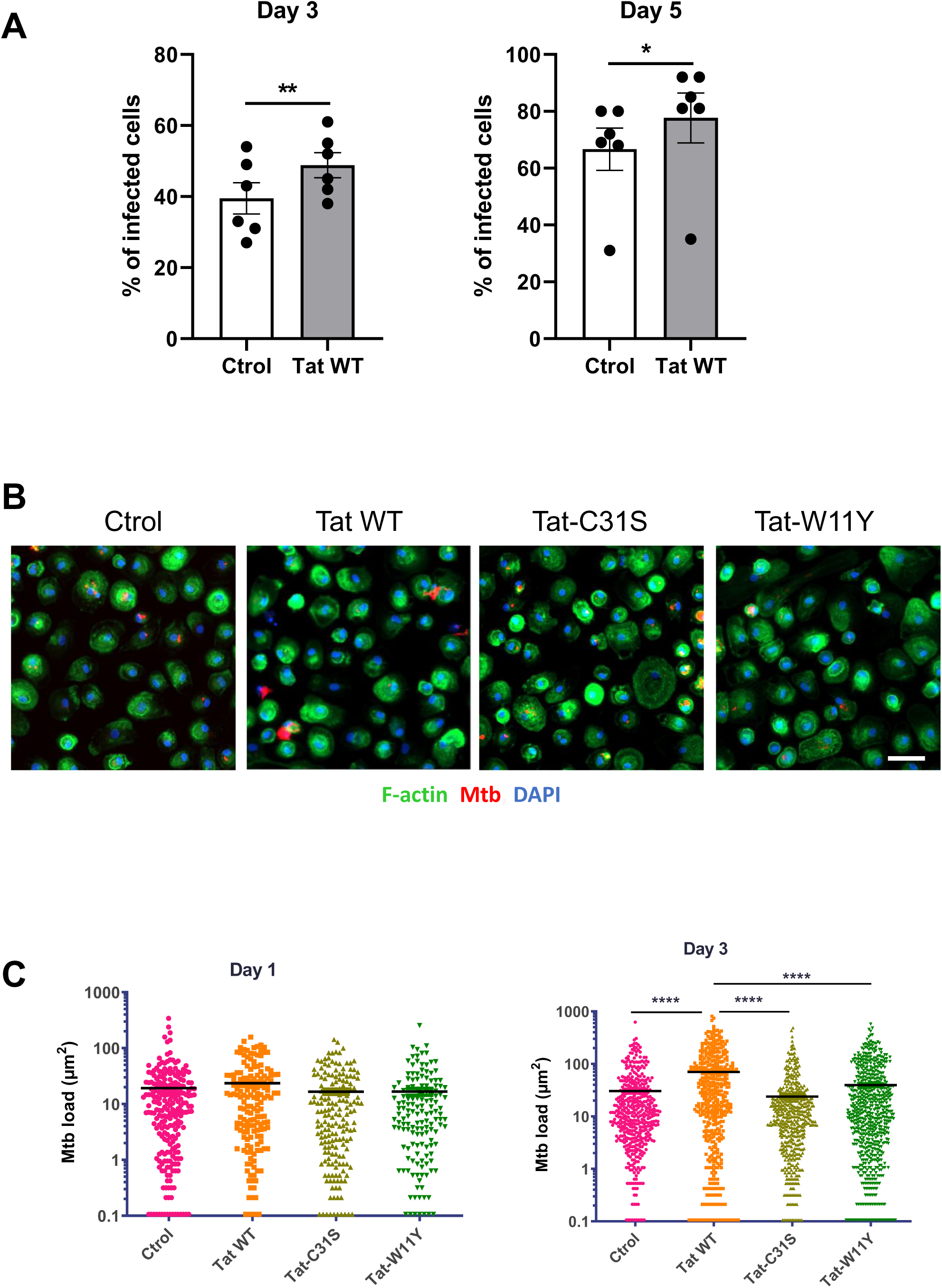
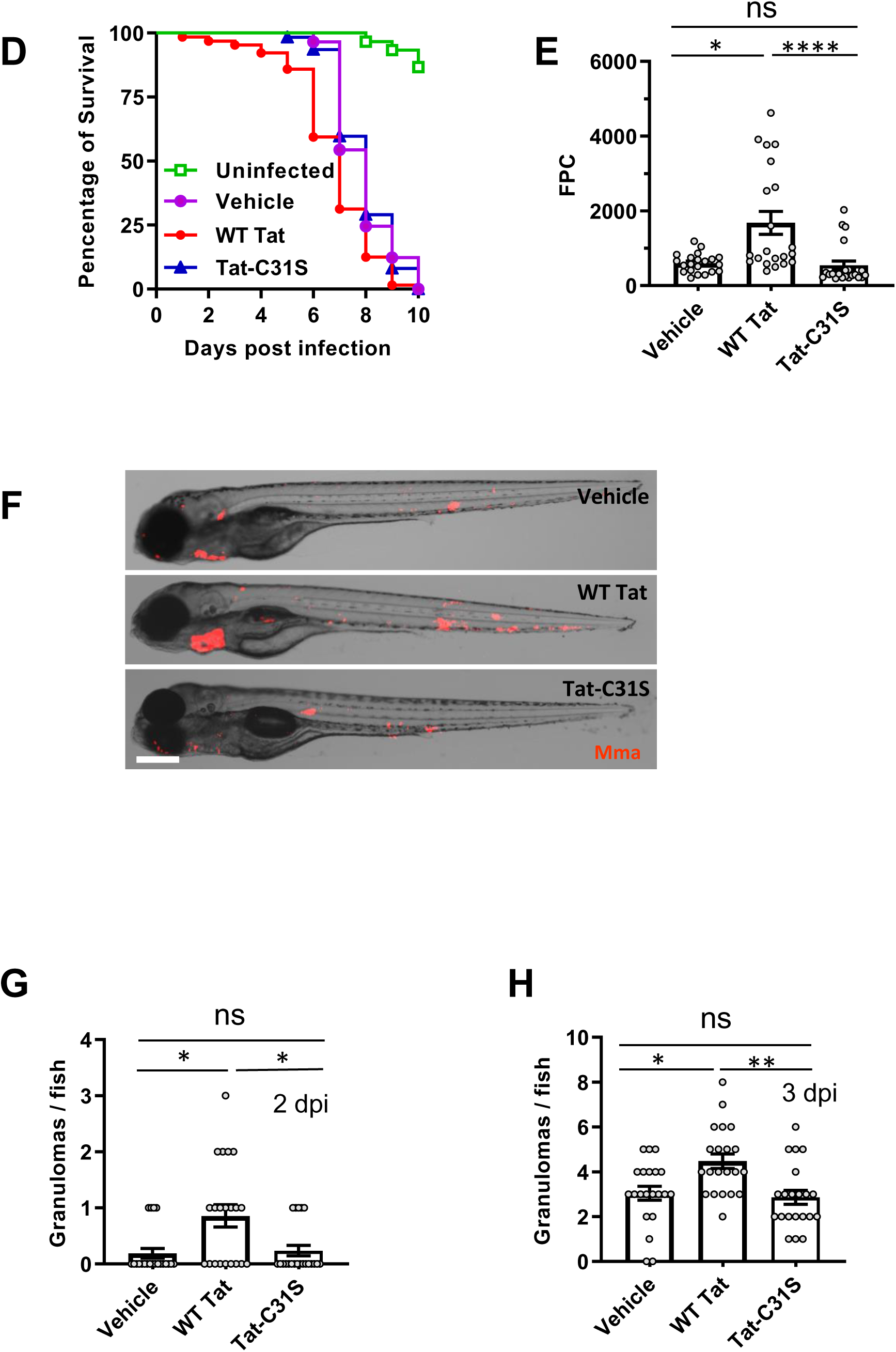
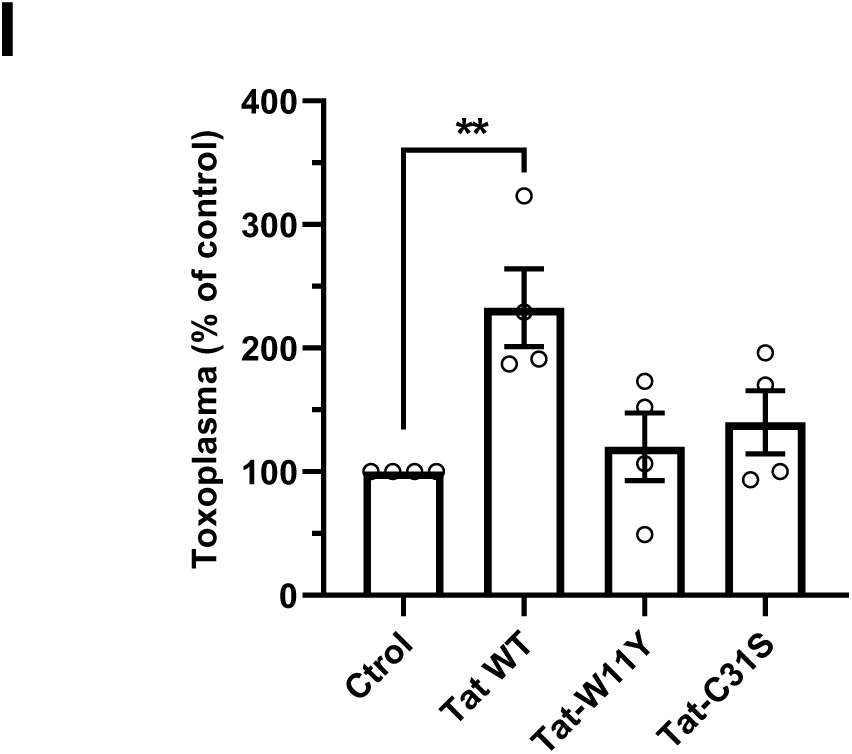
Tat favors the intracellular multiplication of mycobacteria and Toxoplasma. (A-C) Tat favors the multiplication of Mycobacterium tuberculosis in hMDMs. Macrophages were treated with 15 nM Tat for 4 h before adding *Mycobacterium tuberculosis* expressing dsRed (MOI=0.3). The indicated version of Tat (15 nM) was added at day 1 post infection. (A) The percentage of infected cells was determined at day 1, 3 and 5 using bacteria fluorescence. Data are mean ± SEM (n=6 different donors). Paired Student’s t-test (**, p<0.01) and Wilcoxon test (*, p<0.05). (B) Infected macrophages were fixed at day 3 before staining F-actin and DNA using phalloidin and DAPI, respectively. Bar, 20 µm. (C) Mtb intracellular multiplication was monitored using the area of fluorescence spots (semi-automatic quantification). Kruskal-Wallis test compared to control (*, p<0.05; ****, p<0.0001). (D) Zebrafish embryos (n=20-25 for each group) at 24 hpf were infected with tdTomato *Mycobacterium marinum* (Mma) then injected with Tat (∼100 nM final concentration). Tat injection (WT or C31S as a negative control) was repeated at dpi 2 and 3. Survival curves of the Tat groups (WT/ C31S) were compared using Log-rank Test. Vehicle *vs* Tat WT, ***, p<0.001; Tat WT vs Tat-C31S ****, p<0.0001. The graph for Tat-W11Y is in Figure S4A. (E) Bacterial loads per embryo were measured using fluorescence pixel counts (FPC) at 3 dpi. (F) Embryos treated with Tat, WT or C31S were imaged at 3 dpi. Large red spots correspond to granulomas (Figure S5). Bar, 200 µm. (G and H) granulomas were counted at 2 dpi (G) and 3 dpi (H). Mean granuloma numbers at 2 dpi were 0.19 (vehicle), 0.86 (Tat WT) and 0.238 (Tat-C31S), respectively. Kruskal-Wallis test, *, p<0.05; **, p<0.01. (I) hMDMs were treated with 15 nM Tat for 4 h before adding IgG-opsonized *T.gondii* expressing EGFP (MOI=10) for 45 min. Cells were then washed and EGFP fluorescence was read after 24 h using a plate reader. Data are mean ± SEM (n=4 different donors). One-way ANOVA (**, p<0.01).

To confirm these findings using an *in vivo* model, we used zebrafish embryos 24 hours post fertilization (hpf) [36]. Tat injection in the caudal vein did not perturb embryo development or survival (Figure S3). Tat-injected embryos were then infected with *Mycobacterium marinum*, a pathogen that is phylogenetically related to Mtb [37] and for which the zebrafish is a natural host. Survival plots showed that embryos injected with WT Tat were sensitized to infection compared to those injected with the Tat-C31S (Figure 1D) or Tat-W11Y (Figure S4A) negative controls. We also observed that bacterial loads were increased by Tat but not by Tat-C31S (Figure 1E). Tat thus favors mycobacterial multiplication *in vivo*. Infection of zebrafish embryos with *M. marinum* is known to cause the aggregation of macrophages into granuloma-like structures (Figure S5) that are also a hallmark of human tuberculosis. Both infected and uninfected cells are found in these granulomas [38]. We observed that animals injected with Tat WT but not with the Tat-31S negative control had more granuloma after infection (Figure 1F-H). This difference was especially important (+350%) at 2 dpi (Figure 1G). Similarly, more granulomas were found in animals injected with Tat WT compared to those injected with Tat W11Y, although in that case the difference was more obvious after 4 dpi (Figure S4B).

Tat thus favors mycobacterial multiplication both in a cellular model in which human primary macrophages are infected with Mtb and in zebrafish living larvae infected with *M. marinum*. We examined whether this Tat facilitating effect was restricted to Mycobacteria or also observed using another intracellular pathogen, the parasite *Toxoplasma gondii*. The global worldwide prevalence of *T. gondii* in PLWH is ∼1/3, and toxoplasmosis is one of the most common cerebral infection in PLWH [39]. We used IgG-opsonized parasites, because IgG antibodies are typically produced during reactivation of latent infection [40], which is usually the cause of symptomatic toxoplasmosis for PLWH [41]. Upon uptake by macrophages most opsonized parasites are destroyed following phagocytosis, but some of them escape degradation and successfully multiply intracellularly [42]. hMDMs were pretreated with Tat before infection with opsonized *T. gondii* expressing EGFP. Parasitic loads were assayed 24 h after, allowing to measure the intracellular multiplication of *T.gondii* that has a generation time of 6-8h and does not exit cells before 36 hpi [43]. Tat increased by over 2-fold the multiplication of *T. gondii*, while control mutants had no significant effect (Figure 1I). The absence of effect of Tat-C31S and Tat-W11Y that transactivate normally but do not affect PI(4,5)P_2_-dependent traffic [27, 28] indicated that Tat facilitating effect on the multiplication of intracellular pathogens likely involves Tat-PI(4,5)P_2_ interaction.

### Tat inhibits the recruitment of the autophagic machinery on the membrane of vacuoles containing incoming pathogens

We then tried to identify how Tat could protect macrophages from incoming pathogens. Since autophagy was found to control both Mtb and *T. gondii* infection [10, 44], we first examined whether Tat could affect the recruitment of LC3B, a key autophagic marker and player [45] on vacuoles containing incoming opsonized EGFP-expressing *T. gondii*. We observed that, in hMDMs, ∼70 % of vacuoles containing opsonized *T. gondii* were positive for LC3B after 40 min of uptake (Figure 2A-B). This recruitment of LC3B on parasitophorous vacuoles (PVs) was inhibited by ∼50 % when macrophages were pretreated with Tat. Control Tat mutants (Tat-W11Y and Tat-C31S) had no significant effect. This inhibition by Tat of LC3B recruitment on PVs is consistent with Tat facilitating effect on parasite multiplication (Figure 1I). The recruitment on PVs of the autophagy receptor protein p62/SQTM1 (p62) that binds to LC3 was similarly inhibited by Tat but not by Tat-W11Y or Tat-C31S (Figure S6).

**Figure 2.**
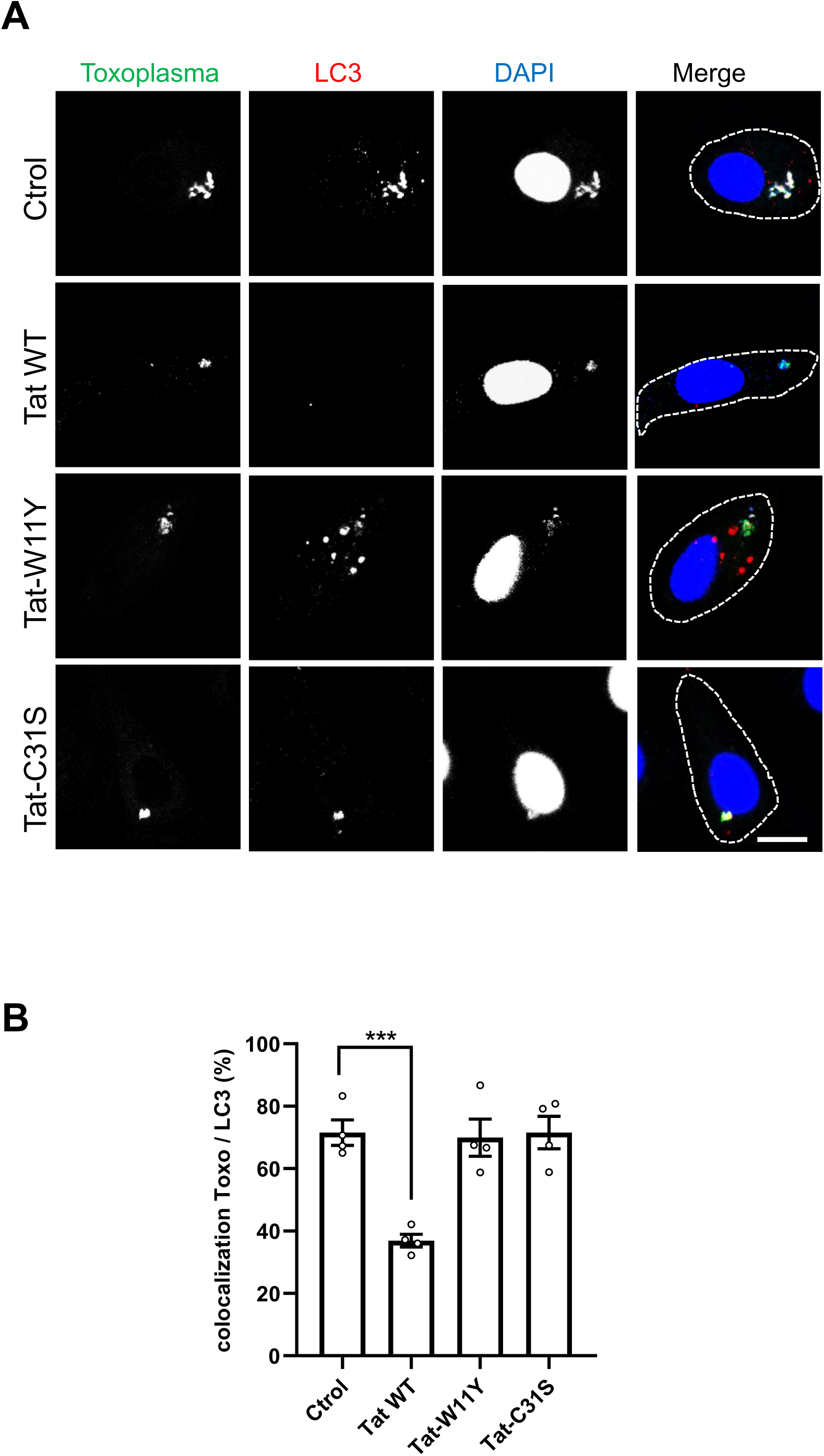
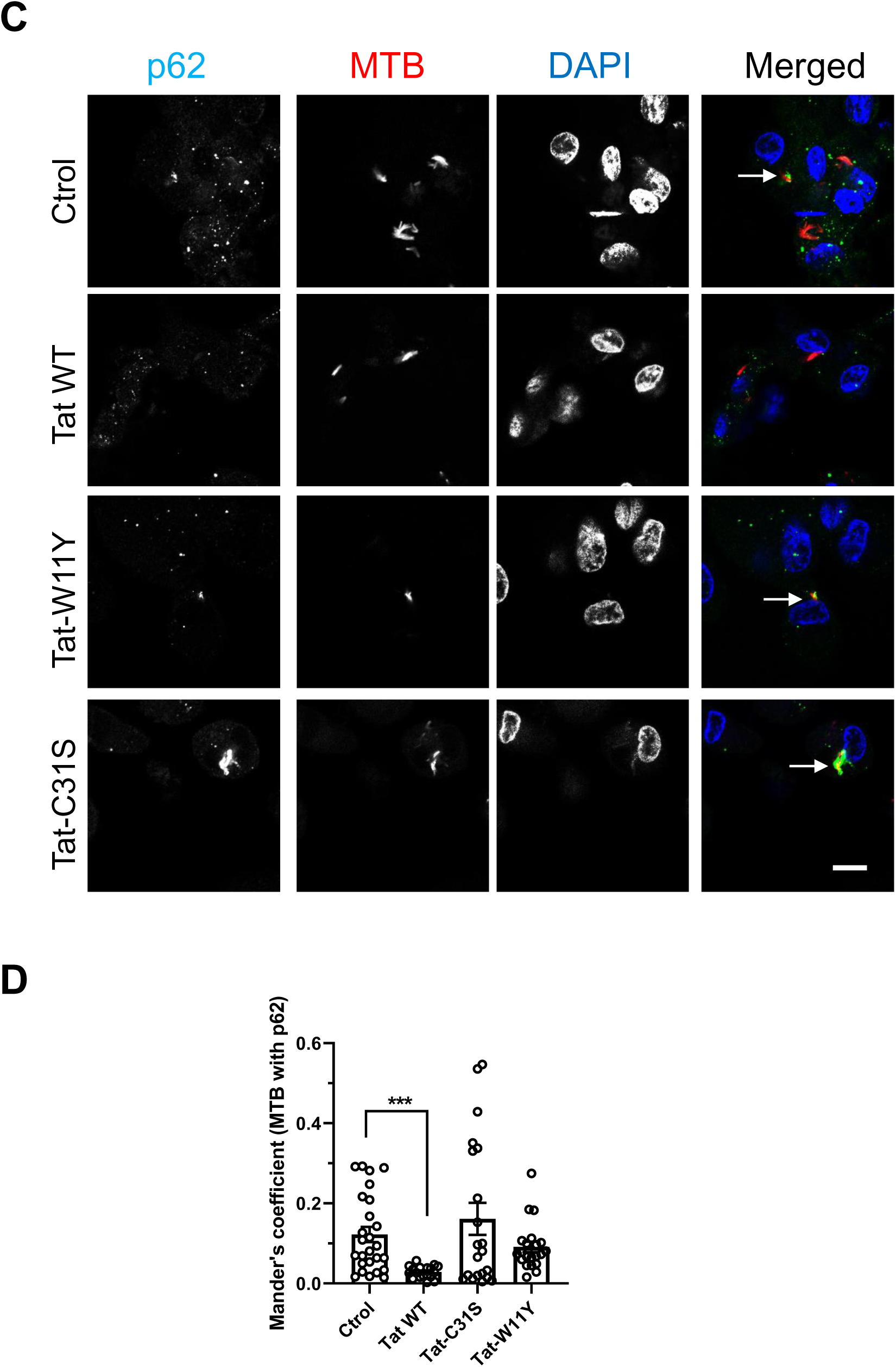
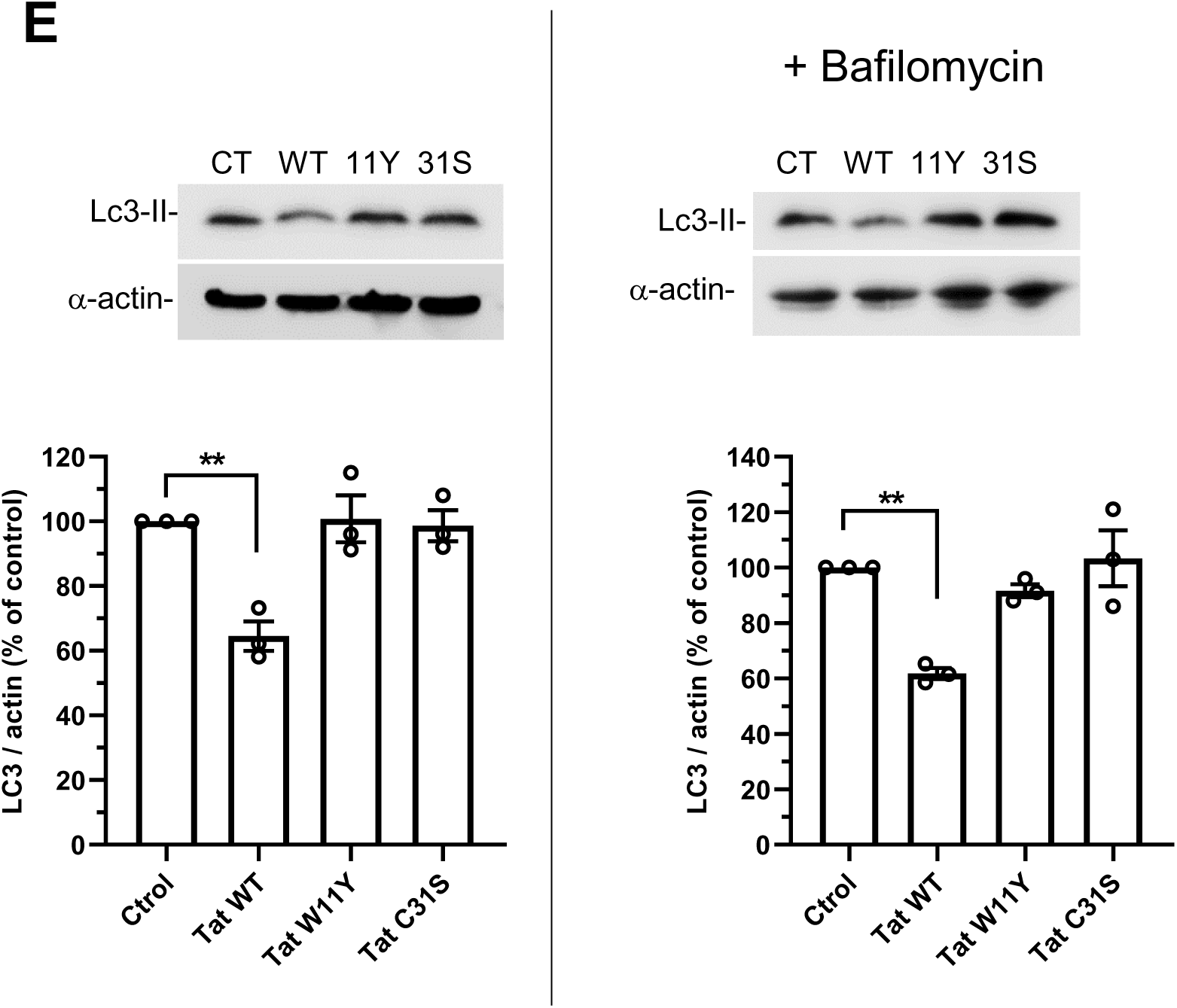
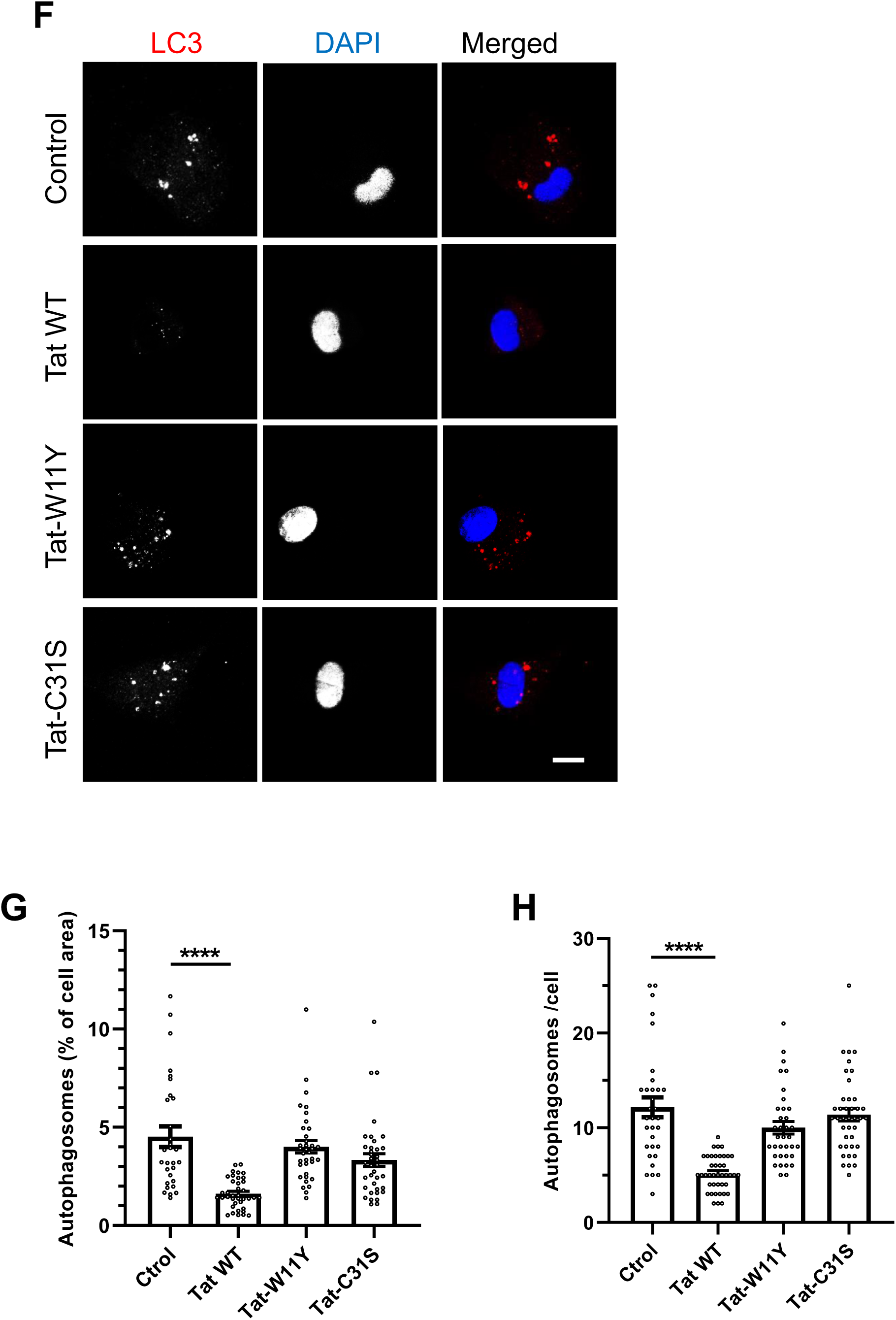
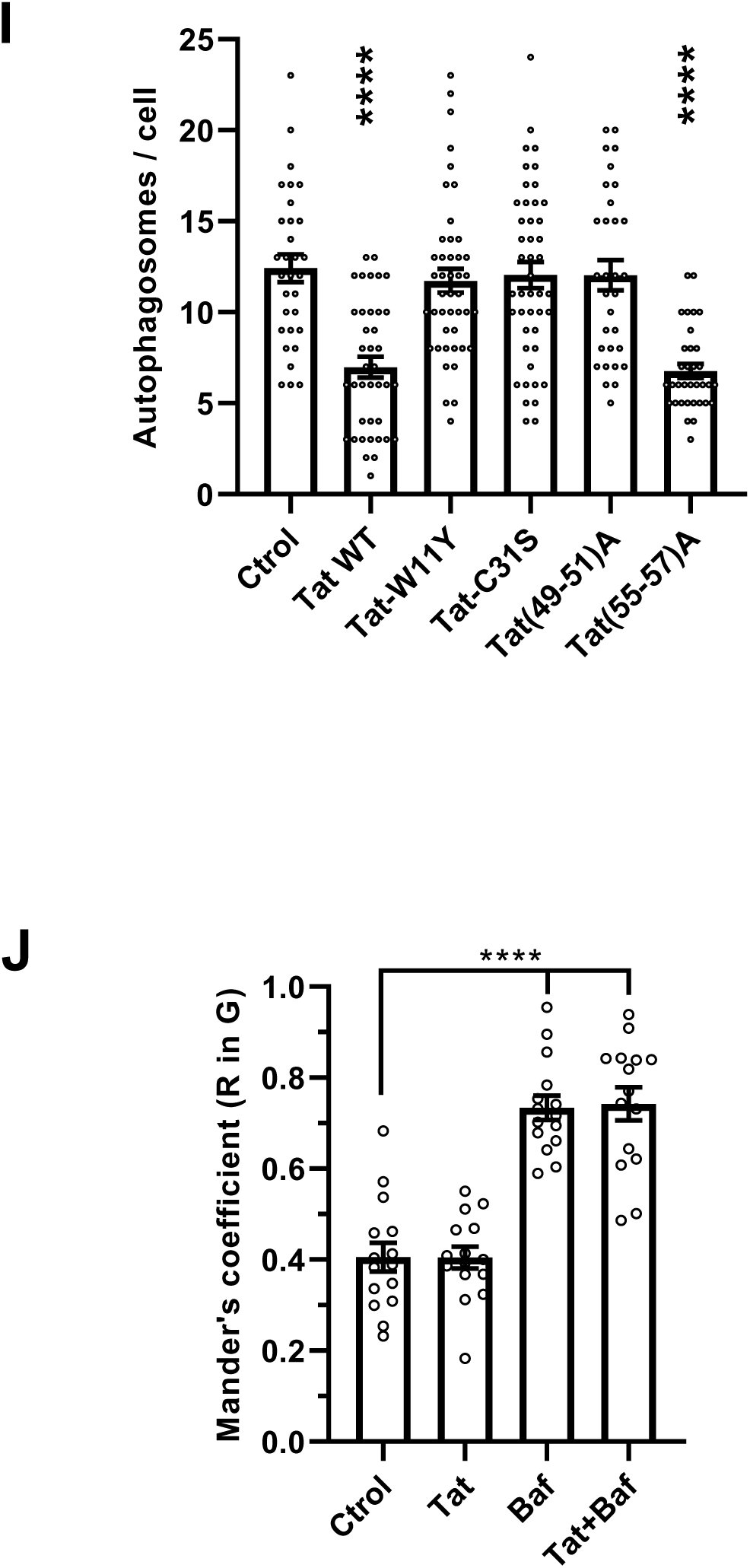
Tat inhibits autophagy. (A) Tat impairs the recruitment of LC3-II on vacuoles containing opsonized *T.gondii*. hMDMs were treated with 15 nM of the indicated version of Tat for 4 h before adding EGFP-expressing opsonized T. gondii (MOI=10). After 40 min, cells were fixed, stained for LC3 and observed using a confocal microscope. Representative optical sections. Bar, 10 µm. (B) Colocalization between intracellular parasites and LC3. At least 30 cells per condition were counted using Fiji for 4 donors. Means ± SEM. One-way ANOVA (***, p<0.001). (C) hMDMs were treated with 15 nM of the indicated version of Tat, then infected with dsRed-expressing Mtb. After 2 days, cells were fixed then stained for p62 before confocal imaging. Arrows point to p62-labelled mycobacteria. Bar, 10 µm. (D) The fraction of Mtb vacuoles labelled with p62 was assessed using Mander’s coefficient (350<n<500 cells per condition, infection rate was 28-35%). Means ± SEM. Kruskal-Wallis test (***, p<0.001). (E) hMDMs were treated with Tat for 4 h. When indicated, 20 nM bafilomycin A1 was then added for 1 h. Cells were harvested and analyzed by anti-LC3 and α-actin western blot. LC3-II only is visible in macrophages using this antibody. The graphs show the quantification from n=3 different donors, setting the control value to 100%. Means ± SEM. One-way ANOVA compared to no-Tat conditions (**, p<0.01). (F) hMDMs were treated with Tat for 4 h before fixation, anti-LC3 immunofluorescence, DAPI staining and confocal imaging. Bar, 10 µm. (G) LC3 spots (autophagosomes) from 30-40 cells were quantified using Fiji and the ratio (total area of the spots) / cell area. Means ± SEM. Kruskal-Wallis test compared to control conditions (****, p<0.0001). (H) Autophagosomes from 30-40 cells were counted. Kruskal-Wallis test compared to control conditions (****, p<0.0001). Results of a typical experiment that was repeated twice. (I) RAW macrophages were transfected with mCherry-LC3B and the indicated version of Tat. Cells were fixed after 18h and imaged using a confocal microscope, before counting LC3B spots (autophagosomes) of 31-45 cells. Representative images are in Figure S7. One-way ANOVA compared to control conditions (****, p<0.0001). Results of a typical experiment that was repeated twice. (J) RAW macrophages were transfected with dsRed-EGFP-LC3B then treated with 15 nM Tat for 4 h. When indicated, 20 nM Baf was added for 1 h and living cells were imaged at 37°C using a spinning-disk confocal microscope. Representative images are in Figure S8. The localization of red spots in green spots was measured for n>15 cells using Fiji and Mander’s coefficient. Means ± SEM. One-way ANOVA compared to control conditions (****, p<0.0001).

p62 is responsible for the autophagic elimination of mycobacteria in macrophages [46]. When we monitored the recruitment of p62 on MTB-containing vacuoles in hMDMs we observed that p62 localized to some mycobacteria vacuoles (Figure 2 C-D). Colocalization quantification showed that ∼12% of Mtb vacuoles were labelled with p62. This is similar to what was observed before in RAW264.7 mouse macrophages (referred to as RAW macrophages hereafter) for the BCG variant of Mtb [46]. When macrophages were pretreated with Tat WT, the extend of p62 labeling of MTB vacuoles decreased by 4-fold while control mutants did not show any significant effect (Figure 2 C-D). This inhibition by Tat of p62 recruitment on Mtb vacuoles is consistent with Tat facilitating effect on mycobacteria multiplication (Figure 1A-C). Altogether these data indicated that Tat interferes with the recruitment of key autophagic effectors onto vacuoles containing intracellular pathogens. This recruitment was not affected by Tat-C31S, therefore indicating that the inhibition of this recruitment by WT Tat relies of the Tat-PI(4,5)P_2_ interaction.

### Tat inhibits basal autophagy in macrophages

To examine whether Tat inhibits basal autophagy in macrophages, we first followed autophagy by monitoring the levels of LC3-II (the membrane-bound form of LC3B associated with autophagic vesicles) in hMDMs by western blots. Bafilomycin A1 blocks the autophagic flux, prevent LC3B-II degradation and allows the autophagic flux quantification [47]. In hMDMs we observed that Baf increased LC3B level by 2-fold (196 ± 39 %, mean ± SEM, n=8). As reported before [48, 49], we observed that Tat inhibits the accumulation of LC3B-II in hMDMs whether Bafilomycin was present of not (Figure 2E). This result indicates that Tat affects an early stage of autophagy instead of the maturation step. Control mutants Tat-W11Y and Tat-31S had no effect. To confirm these biochemical data, we monitored the size of the autophagosome compartment (% of cell area) and the number of autophagosomes using LC3B immunofluorescence. While WT Tat decreased the size of the autophagosome compartment by ∼70% and the number of autophagosomes/ cell by 60% in hMDMs, control mutants Tat-W11Y and Tat-C31S had no effect (Figure 2 F-H).

The absence of effect on the autophagic machinery of Tat-C31S indicates that Tat binding to PI(4,5)P_2_ is likely involved in Tat inhibition of the autophagic machinery. To confirm this point, we co-transfected RAW mouse macrophages with mcherry-LC3B and different Tat mutants, including Tat (49-51)A that is devoid of Tat-PI(4,5)P_2_ binding motif [24]. These mutants versions of Tat are expressed to similar levels upon transfection in RAW macrophages [30]. Mutants in Tat basic-domain (res 49-57) have to be expressed intracellularly because they are poorly endocytosed, preventing their use as recombinant proteins [27]. Tat (49-51)A did not affect autophagy (Figures 2I and S7). The control mutant Tat (55-57)A has the same number of charges as Tat (49-51)A, while retaining the PI(4,5)P_2_ binding motif. Tat (55-57)A behave as WT Tat, while the negative controls Tat-W11Y and Tat-C31S were inactive. Collectively, these data indicate that Tat inhibits autophagy by binding to PI(4,5)P_2_.

Since Tat effect is also observed when the autophagic flux is blocked by Bafilomycin (Figure 2E), Tat presumably inhibits autophagy at an early stage. To confirm this point, we used tandem GFP-mRFP-LC3B that was expressed in RAW macrophages before adding Tat for 5 h and live confocal microscopy. In green/red merged images yellow spots correspond to vesicles with neutral pH, *i.e.* autophagosomes, while red spots correspond to autolysosomes, where acidic pH quenches the EGFP fluorescence. If autophagic flux is inhibited, the accumulation of yellow puncta can be followed using the colocalization of red in green spots [45]. As a positive control for these experiments, we used Bafilomycin that inhibits autophagosome-lysosome fusion [50]. Bafilomycin increased by ∼80 % the colocalization of red in green spots, indicating an accumulation of autophagosomes, while WT Tat did not show any effect (Figures 2J and S8). These results obtained on live cells therefore confirm biochemical data (Figure 2E) indicating that Tat does not affect the autophagic flux but rather interfere with an early stage of the autophagy pathway.

### Tat inhibits CME by preventing the PI(4,5)P_2_-mediated recruitment of AP-2 at the plasma membrane

Tat can affect the recruitment of some proteins by PI(4,5)P_2_ [30] that is well known to recruit key CME players. Since CME is required for autophagy in HeLa [12] and MEF cells [51], we hypothesized that Tat could inhibit autophagy by perturbating CME. We first examined whether Tat affects CME. When hMDMs were treated with 15 nM Tat, the uptake of transferrin, a conventional CME cargo [52] was inhibited by ∼35 % (Figure 3A). We then examined if CME inhibition can affect autophagy in RAW macrophages. To this end we used an siRNA against the AP-2 µ2 chain [53], since AP-2 is recognized as the main CME adaptor [54]. We first checked whether the AP-2 µ2 siRNA affected the recruitment of AP-2σ-EGFP to the macrophage plasma membrane using TIRF microscopy. Compared to the control siRNA, the AP-2µ2 siRNA strongly inhibited (by 45 %) the recruitment of AP-2α to the plasma membrane of macrophages (Figures 3B and S9). The AP-2µ2 siRNA is therefore able to strongly inhibit AP-2 assembly at the plasma membrane.

**Figure 3.**
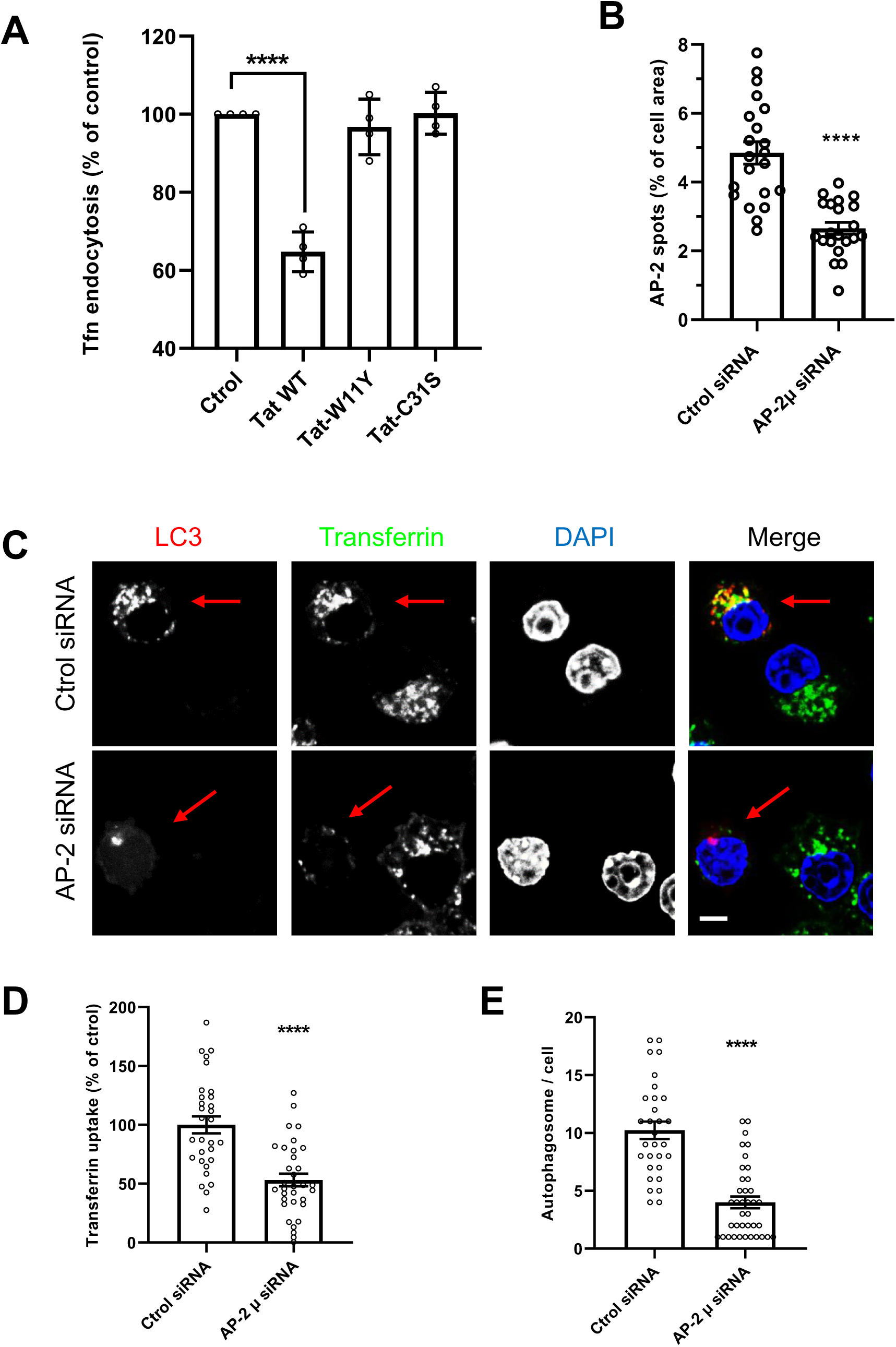
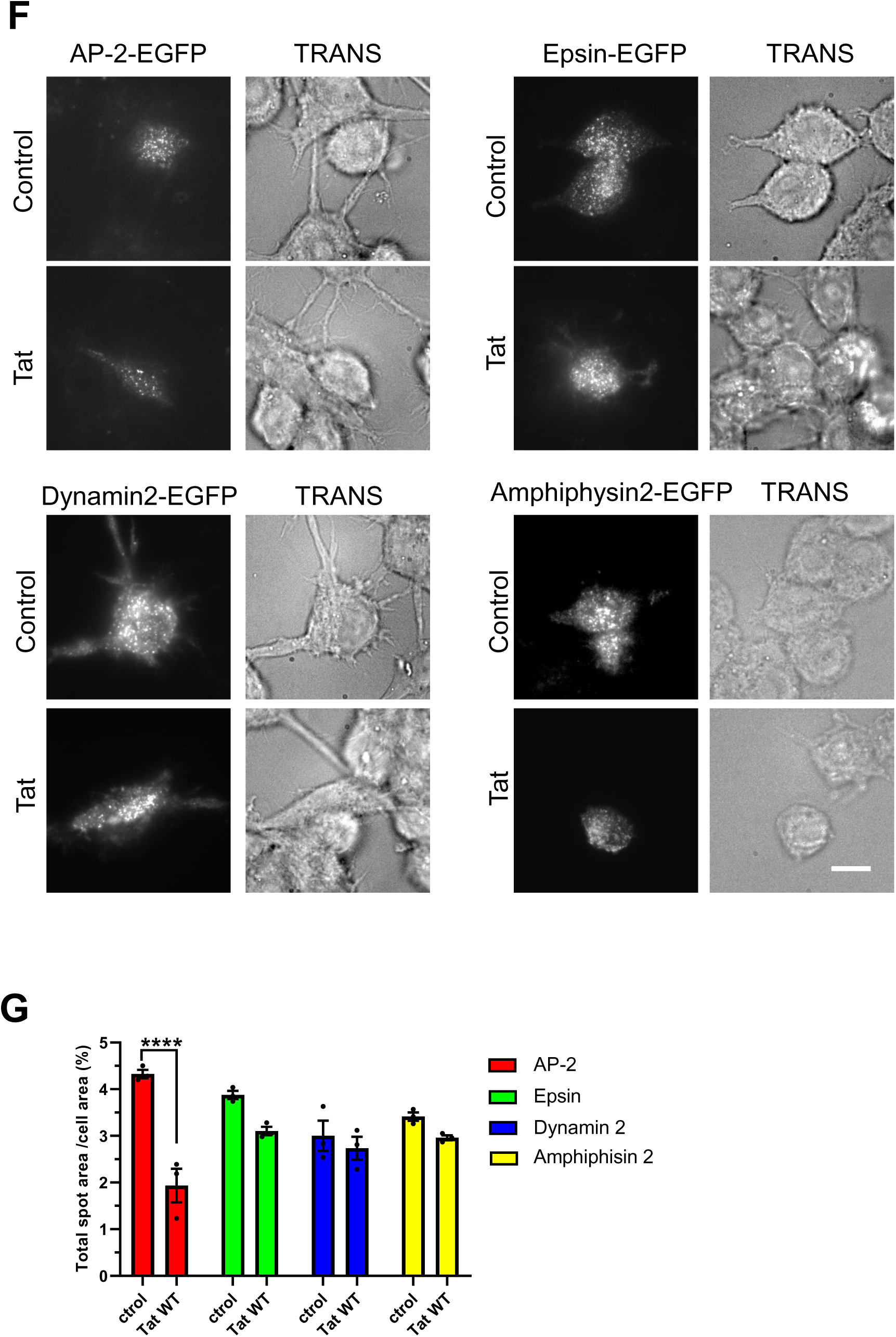
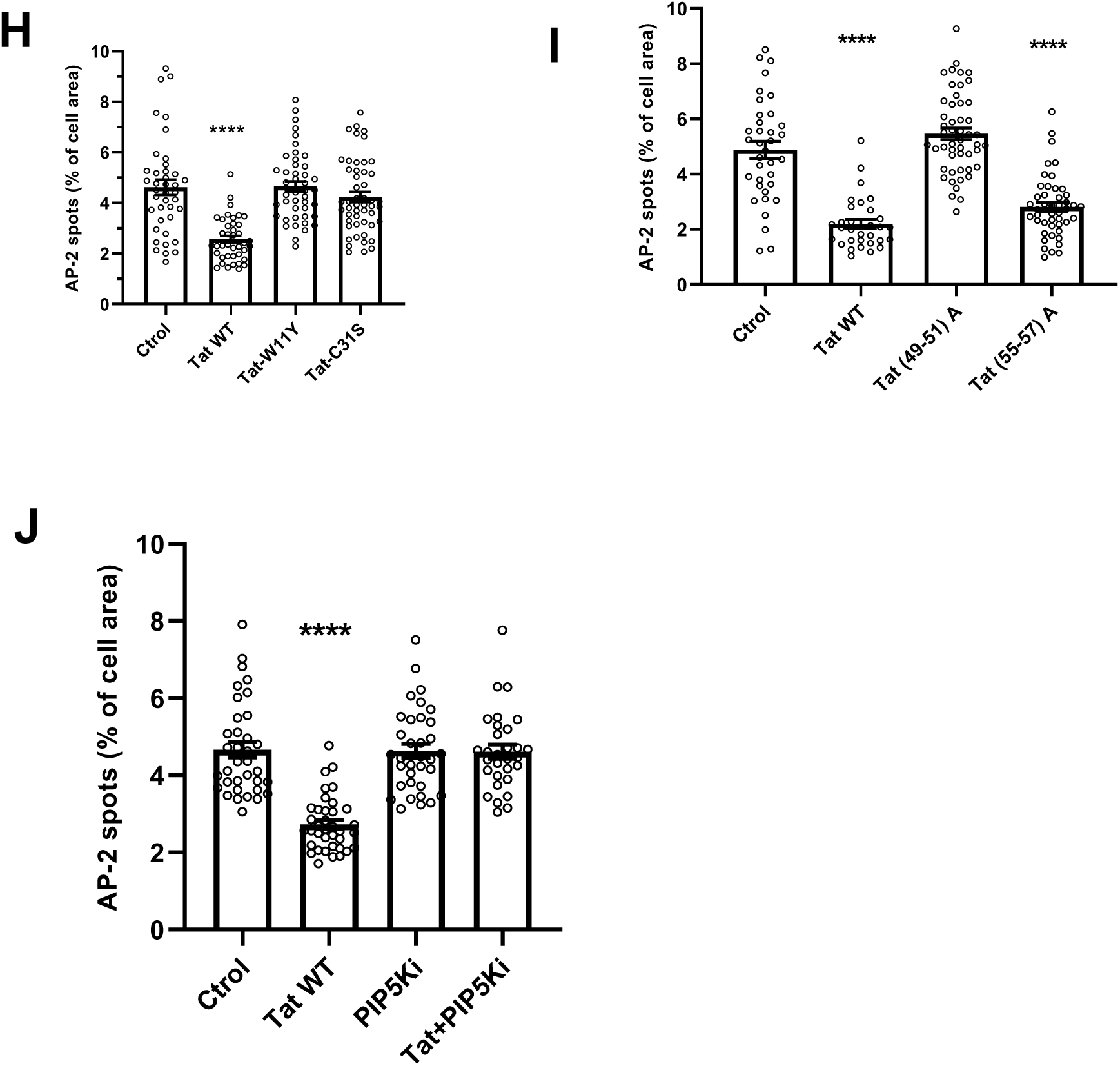
Tat inhibits clathrin mediated endocytosis by preventing the recruitment of AP-2 by PI(4,5)P_2_. (A) Tat inhibits CME in macrophages. hMDMs were treated with 15 nM of the indicated version of Tat for 5 h, then labeled for 5 min with Cy5-transferrin before fixation, imaging using a confocal microscope and quantification of intracellular fluorescence from n=30 cells. Means ± SEM (n=4 different donors). One-way ANOVA (****, p<0.0001). (B) AP-2µ siRNA inhibits AP-2σ2-EGFP recruitment at the plasma membrane. RAW cells were cotransfected with AP-2σ2-EGFP and the indicated siRNA before fixation and imaging by TIRF microscopy. Representative images are in Figure S9. The percentage of cell area occupied by fluorescent spots was quantified using Fiji for n>21 cells. Means ± SEM. Student’s t-test, ****, p<0.0001. (C) RAW macrophages were cotransfected with mCherry-LC3 and the indicated siRNA before labeling cells with Cy5-transferrin for 30 min, fixation, DAPI staining and confocal microscopy. Arrows indicate transfected cells. Bar, 5 µm. (D and E) Transferrin uptake (D) and the number of LC3 spots (autophagosomes, E) were quantified for n=30-35 cells. Means ± SEM. Students’ t-tests, ****, p<0.0001. Results of a typical experiment that was repeated twice. (F) Tat inhibits AP-2 recruitment at the plasma membrane. RAW cells were transfected with the EGFP version of the indicated CME adaptor, treated with 15 nM Tat for 5h before fixation and imaging by TIRF microscopy, using a LED for transmitted light (TRANS). Bar, 10 µm. (G) The percentage of cell area occupied by fluorescent spots was quantified using Fiji for 30 <n< 50 cells. Means ± SEM (n=3 independent experiments). One-way ANOVA (****, p<0.0001). (H) Tat palmitoylation is required for Tat to inhibit AP-2 recruitment. Cells were transfected with AP-2σ2-EGFP, treated with 15 nM of the indicated Tat mutant for 5h before fixation and imaging by TIRF microscopy. Representative images are in Figure S10. The percentage of cell area occupied by fluorescent spots was quantified for 37 <n< 50 cells. Means ± SEM. One-way ANOVA (****, p<0.0001). (I) Tat binding to PI(4,5) is required for Tat to inhibit AP-2 recruitment. Cells were transfected with AP-2σ2-EGFP and the indicated Tat mutant (3/1 ratio), Tat(49-51) A that does not bind PI(4,5)P_2_ or Tat(55-57)A that has the same number of charges but does bind PI(4,5)P_2_. Cells were then fixed and imaged by TIRF microscopy. Representative images are in Figure S11. The percentage of cell area occupied by fluorescent spots was quantified for n> 30 cells. Means ± SEM. Kruskal-Wallis test (****, p<0.0001). (J) Cells were transfected with the AP-2σ2-EGFP vector, a Tat vector and/or a vector expressing the PI(4,5)2 synthesizing enzyme PIP 5-kinase (3/1 ratio). Cells were then fixed and imaged by TIRF microscopy. Representative images are in Figure S12. The percentage of cell area occupied by fluorescent spots was quantified for n> 30 cells. Means ± SEM. Kruskal-Wallis test (****, p<0.0001). Results of typical experiments that were repeated twice.

We then tested its effect on transferrin endocytosis and autophagosome formation, which was monitored using LC3B-mcherry that was co-transfected with the siRNAs. In agreement with its effect on AP-2 recruitment (Figure 3B), the AP-2µ siRNA inhibited transferrin endocytosis by ∼50% and the formation of autophagosomes by ∼60 % (Figure 3C-E). These results are in agreement with those obtained earlier in other cell types [12, 51], and confirm that CME inhibition strongly inhibits autophagy in macrophages. Autophagy inhibition by Tat could thus be due to its capacity to interfere with CME, an effect that likely results from Tat ability to bind PI(4,5)P_2_. We thus tried to identify Tat target(s) in the CME machinery. Among the large number of CME adaptors recruited by PI(4,5)P_2_ at the plasma membrane [55], AP-2 appeared as a strong candidate because it is the main adaptor and requires four PI(4,5)P_2_ molecules to assemble and function properly at the plasma membrane [54]. We used validated EGFP-tagged version of various adaptors [56] transfected into RAW macrophages to examine by TIRF microscopy whether Tat affected their recruitment at the plasma membrane. The recruitment of AP-2 was significantly affected when macrophages were treated with recombinant Tat, while recruitment of the other adaptors epsin and amphiphisin-2, although slightly diminished, was not significantly affected (Figure 3F-G). Dynamin is thought to be recruited at the neck of clathrin-coated pits by PI(3,4)P_2_ [57], and Tat does not bind to PI(3,4)P_2_ [24]. In line with these observations, WT Tat did not affect the recruitment of dynamin at the plasma membrane. Neither Tat-W11Y nor Tat-C31S affected AP-2 recruitment at the plasma membrane (Figure 3H and S10). This result indicated that Tat stable binding to PI(4,5)P_2_ and palmitoylation are required for Tat to perturb AP-2 recruitment. To confirm this result we used Tat-(49-51)A that does not bind PI(4,5)P_2_. Tat (49-51)A did not affect AP-2 recruitment while its control mutant Tat-(55-57)A behaved as WT Tat (Figure 3I and S11). We then tested whether enhancing PI(4,5)P_2_ production at the plasma membrane could prevent Tat to interfere with AP-2 recruitment. We enhanced PI(4,5)P_2_ level at the plasma membrane by ∼2.5 fold using an overexpression of PIP 5-kinase alpha in RAW macrophages [30]. While PIP 5-kinase overexpression did not affect AP-2 recruitment at the plasma membrane, it prevented Tat from inhibiting this recruitment (Figure 3J and S12). Altogether these results showed that Tat prevents AP-2 recruitment at the plasma membrane by masking PI(4,5)P_2_.

### Tat inhibits CME and autophagy by inhibiting AP-2 recruitment

Tat, especially when palmitoylated, has a very strong affinity for PI(4,5) and this enables Tat to prevent the recruitment by this phosphoinositide of cell proteins involved in several machineries involving PI(4,5)P_2_-mediated recruitment [28]. This is for instance the case of Cdc42 whose recruitment is impaired by Tat, thereby inhibiting mannose- and Fcγ-receptor-mediated phagocytosis [30]. It was thus important to examine to what extend the effect of Tat on AP-2 recruitment is responsible for autophagy inhibition. If, in addition to AP-2, other targets are involved in Tat-mediated autophagy inhibition then Tat and AP-2 inhibition should show additive inhibitory effects on autophagy.

To examine this point, we first used the AP-2 siRNA that was cotransfected with a Tat-expressing vector. Tat had to be co-transfected for these experiments because it enters cells using CME [26] and it was therefore not possible to use recombinant Tat whose uptake would have been inhibited by the AP-2 siRNA. To facilitate autophagosome detection cells were co-transfected with mCherry-LC3B. The presence of Tat did not induce further inhibition of transferrin endocytosis (-60%) or drop in autophagosome number (-50%) compared to when the siRNA AP-2 was used alone (Figure 4A-B). A Cdc42 siRNA, that efficiency depletes RAW macrophages in Cdc42 (Figure S13) and strongly inhibits FcγR-mediated phagocytosis [30], did not significantly affect neither transferrin uptake nor autophagy (Figure 4A-B). These results therefore indicated that AP-2 is the main Tat target responsible for autophagy inhibition by Tat. To confirm that autophagy inhibition by Tat is essentially due to CME perturbation, we used two CME inhibitors, Pitstop2 and Dyngo 4a that interfere with clathrin and dynamin function, respectively, and strongly inhibit transferrin endocytosis [58, 59]. In RAW macrophages, these molecules inhibited transferrin endocytosis by ∼50% and similarly decreased autophagosome number per cell. Again, no further inhibition was observed when Tat was present (Figure 4C and S14). To inhibit dynamin2 function we also used the dominant negative version (K44A) of dynamin2 that prevents the formation of clathrin-coated vesicles [60]. Whereas no increase in transferrin uptake or autophagosome number was observed upon WT dynamin2 overexpression, expression of dominant negative dynamin2 (K44A) reduced Tf endocytosis and the number of autophagosome by ∼50%. The presence of Tat did not further reduce these values (Figure 4D and S15). Hence, none of the tested CME inhibitors (AP-2 siRNA, drugs or dominant negative dynamin) synergizes with Tat to inhibit autophagy. Altogether these data indicate that the inhibition of autophagy by Tat essentially result from CME perturbation due to interference with AP-2 recruitment by PI(4,5)P_2_. We then used Tat mutants. They are expressed to similar levels when transfected into RAW cells [30]. CME and autophagy were only affected by Tat mutants able to bind and stay on PI(4,5)P_2_, *i.e.* WT Tat and Tat(55-57)A (Figure 4E and S16).

**Figure 4.**
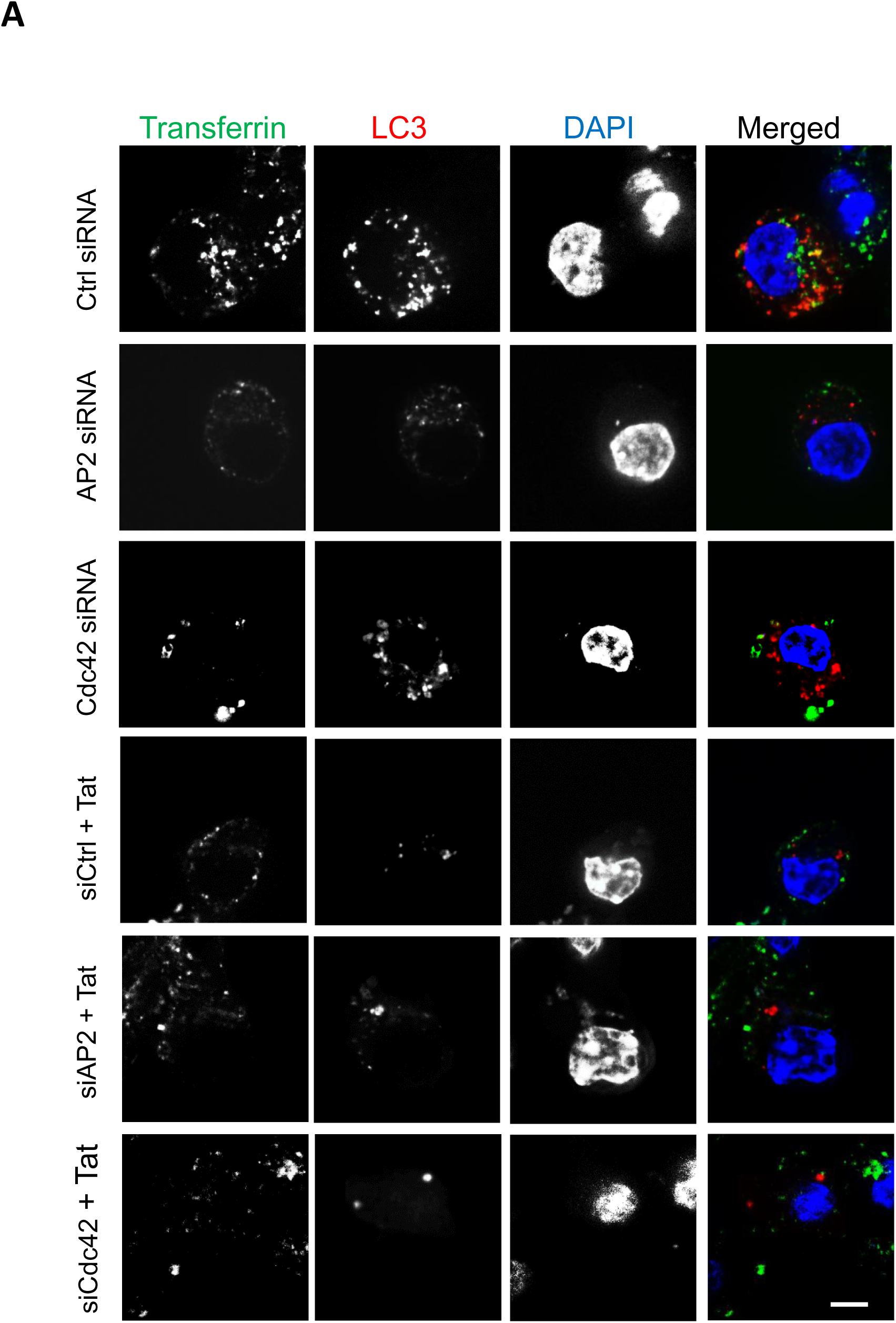
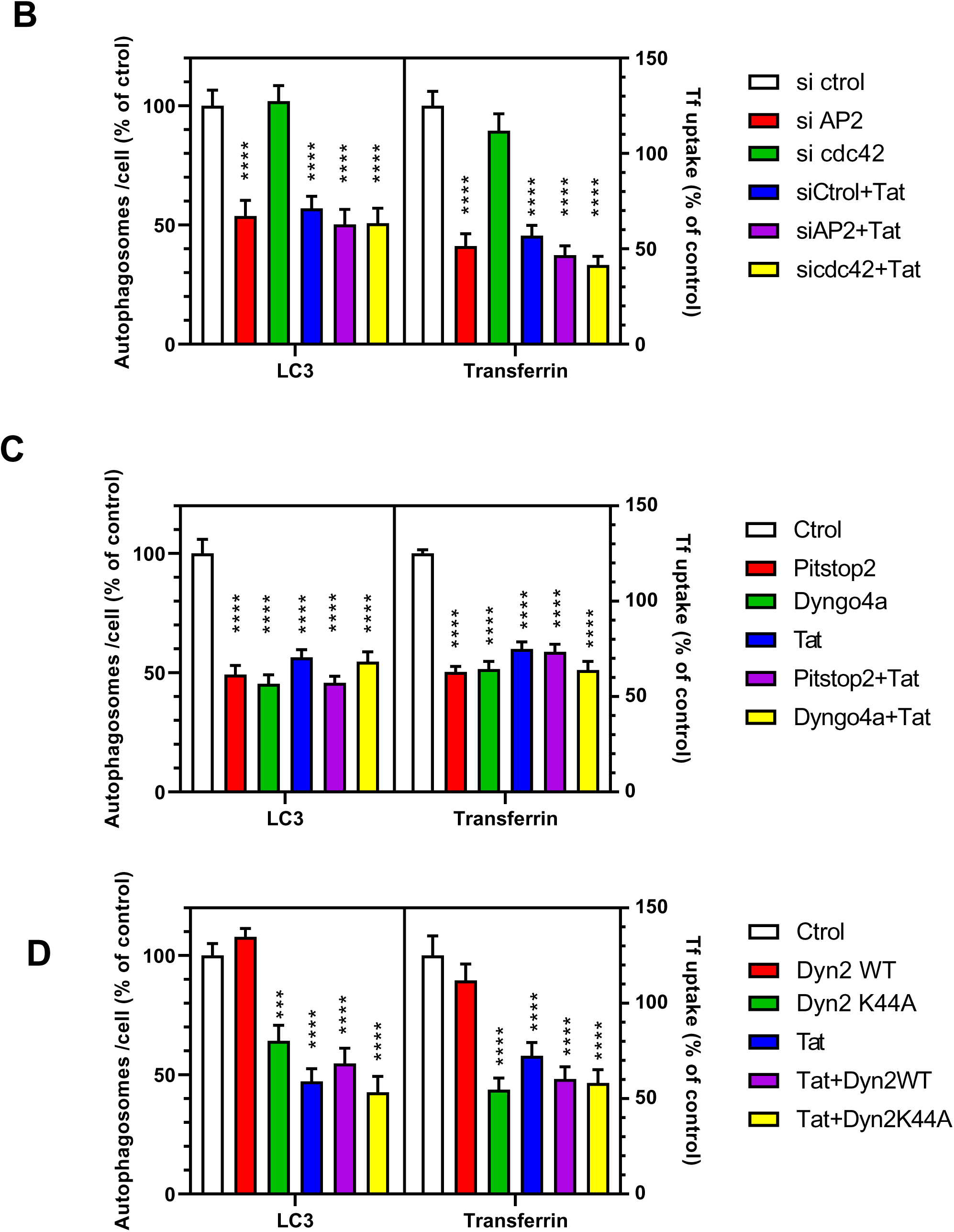
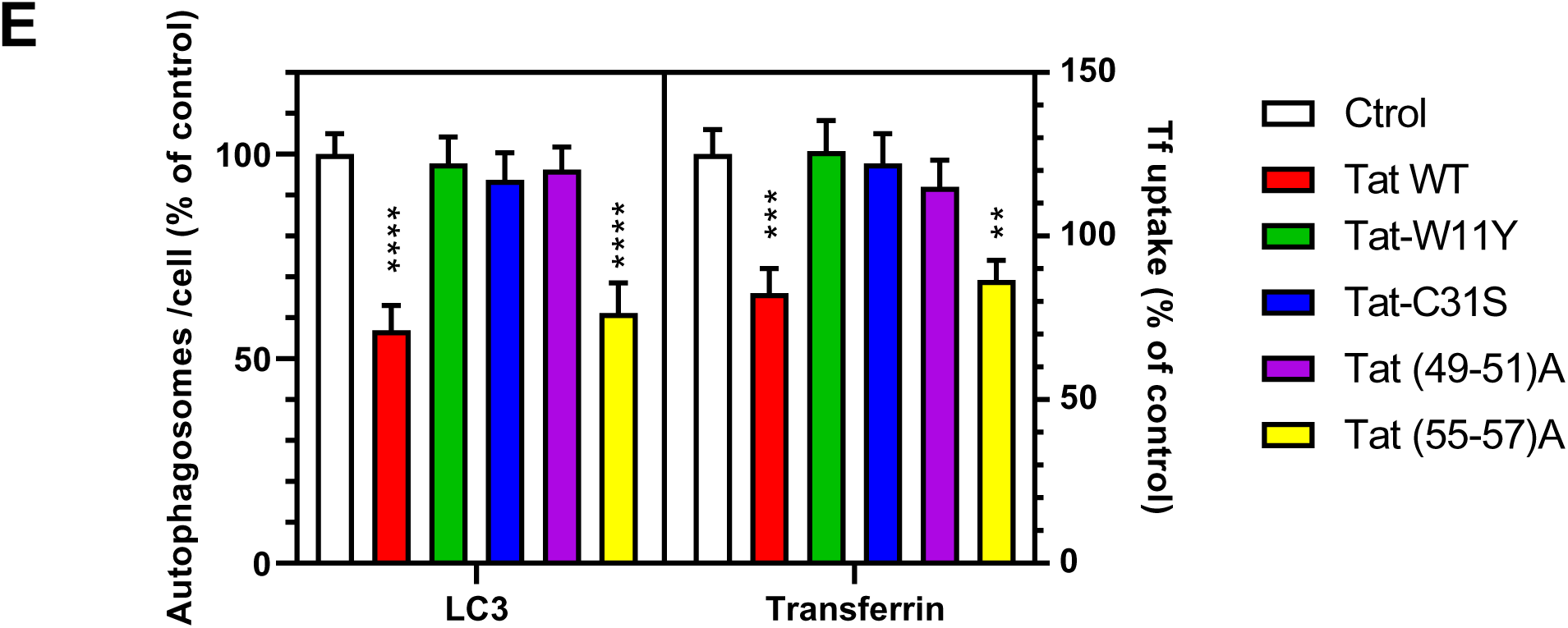
The perturbation of AP-2 recruitment by Tat is responsible for autophagy inhibition. (A) RAW macrophages were cotransfected with mCherry-LC3, Tat and the indicated siRNAs before labeling cells with Cy5-transferrin for 30 min, DAPI staining and confocal microscopy. Bar, 5 µm. (B) Transferrin uptake and the number of LC3 spots (autophagosome) were quantified for n=20-30 cells. (C) RAW cells were transfected with mCherry-LC3 and Tat as indicated, treated with 30 µM Pistop2 or Dyngo4a for 90 min before labeling cells with Cy5-transferrin for 30 min, fixation, and confocal microscopy. Representative images are in Figure S14. Transferrin uptake and the number of LC3 spots (autophagosome) were quantified for n=35-51 cells. (D) RAW cells were co-transfected with mCherry-LC3, dynamin 2 (WT or K44A as indicated) and Tat as indicated. 18h after they were labelled with Cy5-transferrin for 30 min before confocal microscopy. Transferrin uptake and the number of LC3 spots (autophagosome) were quantified for n=35-40 cells. Representative images are in Figure S15. (E) RAW cells were co-transfected with mCherry-LC3 and the indicated Tat mutant. 18h after, cells were labelled with 100 µM Cy5-transferrin for 30 min, before fixation and confocal microscopy. Representative images are in Figure S16. Transferrin uptake and the number of LC3 spots (autophagosome) were quantified for n=35-40 cells. Data are means ± SEM. One-way ANOVA compared to control (**, p< 0.01; ***, p<0.001; ****, p<0.0001). Results of typical experiments that were repeated twice.

### Tat enables the accumulation of lipid droplets

Autophagy can lead to the degradation of intracellular pathogens, but it is also involved in the turnover of lipid droplets (LD), which are used by many intracellular pathogens to obtain lipids that fuel their multiplication, and this is the case for both Mtb and *T. gondii* [61]. We examined whether Tat affects LD concentration in MDMs using BODIPY 493/503 to stain lipid droplets [62]. We found that WT Tat increased by ∼60% the LD content of macrophages, (Figure 5 A-C) while negative controls Tat-W11Y and Tat-C31S had no effect.

**Figure 5.**
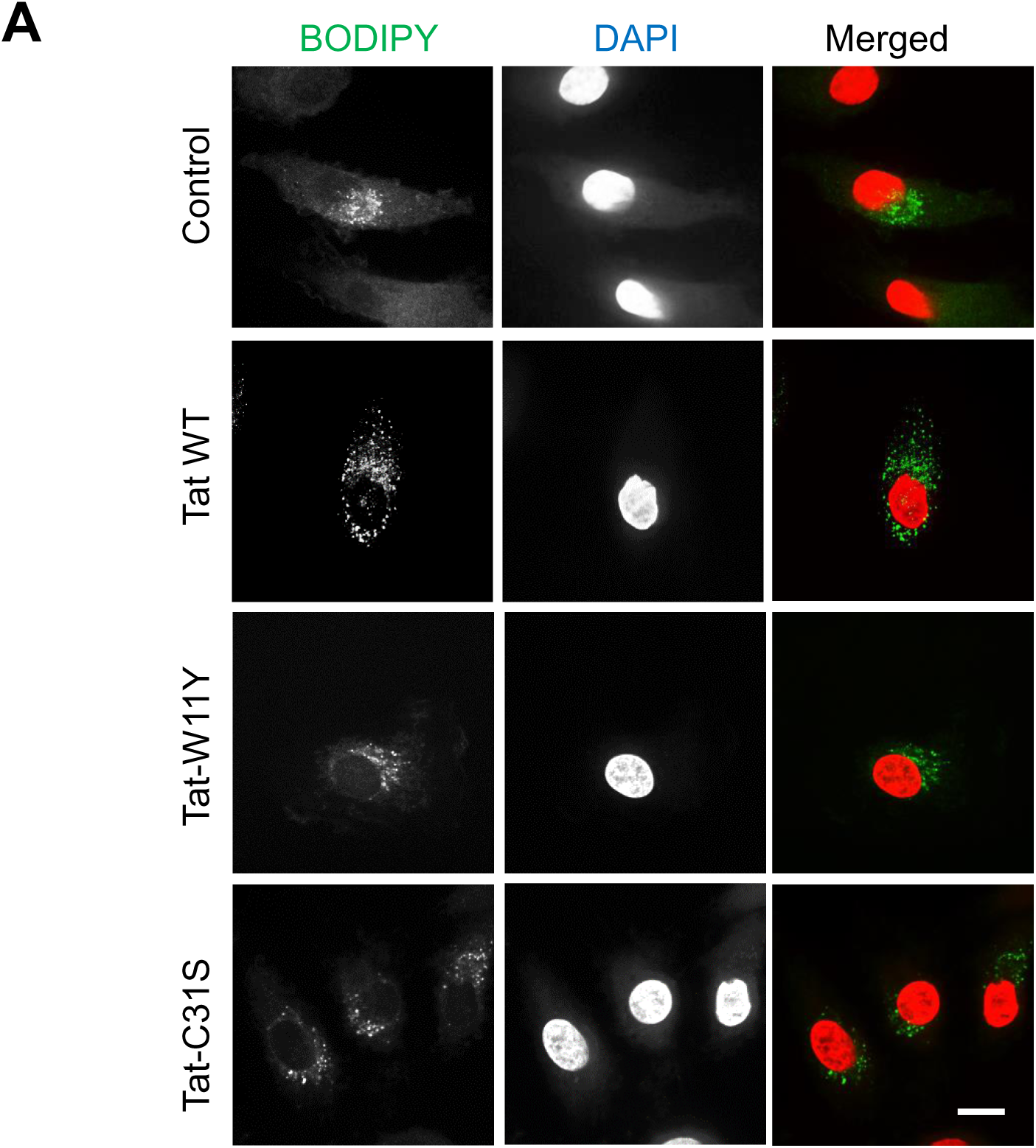
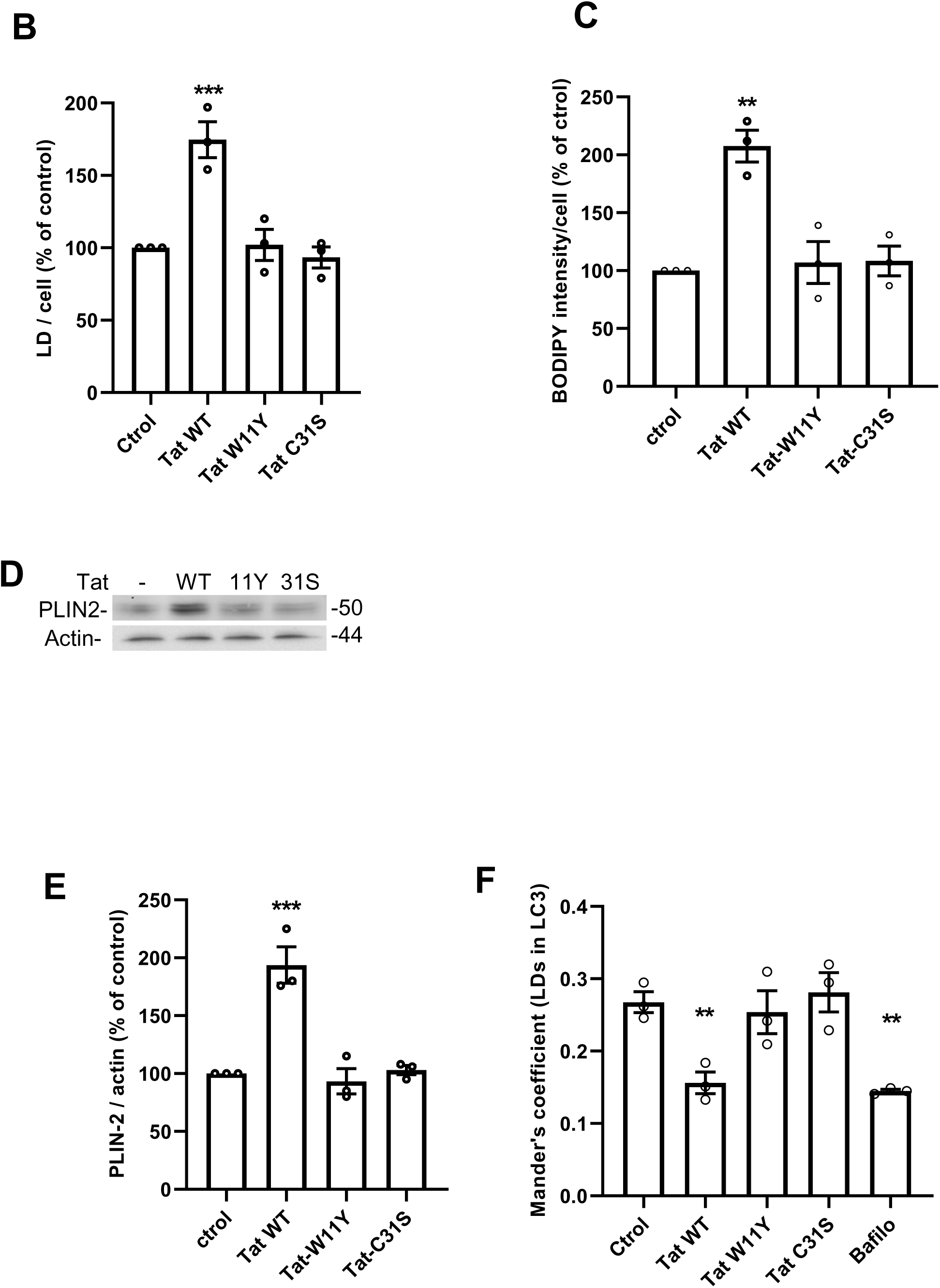
Tat increases the number of lipid droplets in macrophages. (A) hMDMs were treated with 15 nM of the indicated version of Tat for 5 h before fixation, staining lipid droplets with BODIPY 493/503, and imaging using a confocal microscope. Bar, 10 µm. (B) LDs were counted for 30 < n < 46 cells. (C) The intensity of BODIPY staining per cell was measured using Fiji. (D) After Tat treatment, macrophages were scraped for anti-PLIN2 and anti-actin Western blots. (E) Western blot bands were quantified and normalized to control values. Data are means ± SEM from n=3 donors. One-way ANOVA (**, p<0.01; ***, p<0.001). (F) RAW cells were transfected with mCherry-LC3, treated with 10 nM of the indicated Tat mutant for 5 h or 100 nM bafilomycin A1 (Bafilo) for 2 h before staining with BODIPY 493/503, fixation and imaging by confocal microscopy. Representative images are in Figure S17. Localization of lipid droplets in LC3^+^ structures was quantified using Mander’s coefficient. Mean ± SEM of 3 independent experiments. One-way ANOVA compared to control (**, p< 0.01).

To confirm these findings, we monitored the level of the LD-binding protein adipophilin/ perilipin 2 (PLIN2) whose level correlates to macrophage LD content [15]. We found that Tat increased the level of PLIN2 by ∼90 %, while negative Tat controls had no effect (Figure 5 D-E). Hence Tat increases the number of LDs in primary macrophages. Autophagy is known to regulate regulates LD concentration and, in mouse primary macrophages, ∼ 60 % of LDs are positives for LC3B [15]. Using mCherry-LC3B transfected RAW macrophages we found that ∼30% of LDs colocalize with autophagosomes (Figure 5F and S17). Tat decreases this colocalization by ∼40%, while control Tat mutants had no effect. In these experiments, Tat was as efficient as the autophagic inhibitor Bafilomycin. These results are in agreement with the inhibitory effect of Tat on autophagy (Figure 2 and 4).

## Discussion

Despite their crucial epidemiologic relevance, comorbidities such as HIV-1/Mtb synergy remain poorly characterized at the fundamental level [6]. HIV-Tat was detected at nanomolar levels in the sera and cerebrospinal fluid of ART-treated, virally-suppressed PLWH [22, 23, 25]. Here we used Mtb primary target cell, the macrophage, to investigate whether exogenous HIV-Tat could affect Mtb multiplication. We observed that Tat enhanced Mtb multiplication in hMDMs (Figure 1A-C). These results were confirmed *in vivo* when zebrafish larvae were infected with *M. marinum* (Figure 1D-G). These findings are consistent with a previous report indicating that Tat-transgenic mice are more susceptible to Mtb infection [63]. Moreover, Tat was found to facilitate the multiplication of opsonized *T. gondii,* indicating that HIV Tat could promote the multiplication of several intracellular pathogens. Since Tat facilitating effect relies on autophagy inhibition, Tat likely facilitates the multiplication of pathogens that are effectively targeted and destroyed by autophagy. Conversely, Tat might not inhibit the multiplication of pathogens that subvert and hijack the autophagy pathway for their benefit [64].

Tat was previously shown to perturb the autophagic pathway in macrophages [48, 49], but the underlying mechanism responsible for autophagy inhibition remained unknown. We showed here that, in macrophages, Tat inhibits the PI(4,5)P_2_-mediated recruitment of AP-2 the main CME adaptor, thereby inhibiting CME and, in turn, autophagy. Tat mutants unable to bind PI(4,5)P_2_ or be palmitoylated such as Tat-W11Y, Tat-C31S and Tat(49-51)A failed to affect autophagy. This partial (∼50%) inactivation of the autophagic pathway by Tat likely favors the multiplication of intracellular pathogens both directly by preventing their degradation, and indirectly by enabling lipid droplet accumulation. This accumulation of neutral lipids is indeed known to facilitate the multiplication of pathogens by increasing the size of the macrophage lipid storage compartment, thereby enabling to fuel pathogen replication [65].

Regarding the regulation of the entry step of Mtb and IgG-opsonized toxoplasma multiplication by Tat, the mannose receptor has been identified as an important receptor for the phagocytosis of virulent Mtb by macrophages [66], while Fcγ receptors allow the uptake of IgG-opsonized particles [67]. Both receptors require Cdc42 to fulfil their phagocytic function [67, 68]. Because Tat inhibits the PI(4,5)P_2_-mediated recruitment of Cdc42 at the plasma membrane [30], Tat thus likely has a negative effect on the entry of Mtb and opsonised toxoplasma due to the inhibition of the phagocytic activity of the FcγR and the mannose receptor [30] and a positive effect on Mtb and Toxoplasma multiplication due to autophagy inhibition (this study). Nevertheless, because phagocytosis inhibition by Tat is partial (∼45% [30]), it cannot entirely block the uptake of these pathogens by macrophages and, once the pathogens are inside, the positive effect on Mtb and toxoplasma multiplication due to autophagy inhibition is the only effective one and thus becomes dominant.

The facilitating effect of HIV-1 infection on Mtb multiplication is multifactorial[5, 6]. The implication of Tat-induced autophagy inhibition in tuberculosis development in PLWH is thus difficult to pinpoint. Nevertheless, it is interesting to note that in India and South East Asia most infections result from an HIV-1 subtype C that has a non-palmitoylable Tat-S31 [69]. When the cumulative Mtb-associated deaths in the HIV-negative population was examined on the 2010-2023 period it was observed that Mtb-associated deaths is 3-fold higher in South east Asia than in Africa (19 vs 5.9 millions) while Mtb-associated deaths of PLWH is much higher in Africa compared to India (5.1 vs 0.91 millions) [1]. This enhanced susceptibility of PLWH to Mtb in Africa could be linked to the fact that in this region of the world subtypes with Tat-C31 only are present [69] thereby facilitating Mtb multiplication in PLWH. Moreover, as discussed earlier [28], the Indian subtype C is also much less neurotoxic compared to subtypes encoding for palmitoylable Tat [69, 70]. This could be linked to the inability of unpalmitoylated Tat to remain on PI(4,5)P_2_ and affect neurosecretion [28]. Altogether, epidemiological data of the Indian subtype C and its non-palmitoylable Tat indicate that circulating Tat could be involved in the development of both neurological disorders and Mtb that affect PLWH [6, 71].

We here described a mechanism enabling HIV to favor Mtb multiplication due to the action of secreted Tat that is present in the sera and CSF of PLWH despite ART [22, 23, 25]. This effect of Tat is consistent with the observation that PLWH under ART still remain 4-to 7-fold more sensitive to Mtb infection than the HIV-negative population [2, 4].

## Materials and Methods

### Chemicals

Most chemicals were obtained from Sigma. CME inhibitors were from Cayman Chemical company and BODIPY 493/503 from Thermo scientific.

### Protein, antibodies and expression vectors

Tat (HXB2, 86 residues) WT and mutants were prepared as described [28]. Fluorescent transferrin was prepared using a Cy5-labeling kit as recommended by the manufacturer (GE Healthcare). Anti-LC3 was from Nanotools (# LC3-5F10), anti-perilipin-2 for Western blots was from Progen (GP41), anti-β actin (sc-47778) and anti-p62 (sc-28359) from SantaCruz. Secondary antibodies (peroxidase or fluorochrome labeled) were from Jackson Immunoresearch except anti-guinea pig-IR dye 800 that was from Licor (#925-32411).

Tat expression vectors have been described [28]. Vectors expressing EGFP-tagged version of AP-2σ2 (Addgene # 53610), epsin, dynamin 2ab and amphiphisin 2 have been validated [56]. The increase of PI(4,5)P_2_ induced by PIPKiα overexpression in RAW cells was shown before [30]. The pCR3.1 vector expressing dynamin 2ab and its dominant negative version (K44A) was a kind gift of David McNiven and Nathalie Sauvonnet.

### Cell culture

For the preparation of human monocytes, freshly drawn human blood was obtained from the local blood bank (Etablissement Français du Sang, Montpellier, agreement # 21PLER2019-0106). Peripheral blood mononuclear cells were prepared by density gradient separation on Ficoll–Hypaque (Eurobio) before isolating monocytes using CD14 microbeads (Miltenyi Biotec’s). Monocytes were then differentiated in macrophages for 6–8 days in RPMI supplemented with 10 % FCS and 50 ng/ml of macrophage colony-stimulating factor (Immunotools).

RAW264.7 mouse macrophages were obtained and cultured according to the recommendations of the American Tissue Culture Collection. They were checked monthly for the absence of mycoplasma contamination and transfected with plasmids using Jet-Optimus (Ozyme, France) as recommended by the manufacturer. For plasmid and siRNA co-transfection, lipofectamine 2000 was used as recommended by the manufacturer. SiRNA against AP-2 was a Smartpool on-target plus AP2m1 (11773; a mix of 4 siRNAs against AP2µ2) from Dharmacon. The control siRNA was a mission esiRNA against Renilla luciferase (EHURLUC). It was from Sigma-Aldrich, as was the Cdc42 siRNA [30].

### Cytokine and NF-κB activation assays

To monitor cytokine secretion hMDMs were treated with 15 nM Tat for 5 h before harvesting cell supernatants and assaying cytokine concentration using a LEGENDplex human CD8/NK kit for IL-6, IL-10 and TNF-a, and a Bio-Techne DuoSet ELISA for IL-1β. Assays were performed in triplicate for n=3 different donors, following the manufacturer instructions.

To assay NF-κB activation, RAW macrophages were transfected with a vector containing the Firefly gene behind 3 NF-κB sites [72] and another with the Renilla gene behind the thymidine kinase promoter (Promega). Cells were treated with 15 nM Tat or 0.5 µg / ml *E. coli* LPS for 5 h. Cells were then lysed for luciferase activities assayed using Dual-Glo luciferase assay system (Promega).

### Infection with Toxoplasma

Tachyzoites of the RH strain of *T. gondii* expressing EGFP [73] were maintained in human foreskin fibroblasts (American Type Culture Collection, CRL 1634). Extracellular tachyzoites were opsonized by incubation in a 1:200 dilution of heat-inactivated Toxoplasma-seropositive rabbit serum (a kind gift of JF Dubremetz [74]) for 30 min at 37 °C. Opsonized parasites were washed twice in HBSS then counted and resuspended in Dulbecco’s modified Eagle’s medium (DMEM) supplemented with 10 mM Hepes before adding to macrophages. hMDMs were pretreated with 15 nM Tat for 5 h before adding EGFP-*T. gondii*. An MOI of 1-3 Toxoplasma / macrophage was used and parasites were first centrifuged on cells (1 min x 20 g) before incubation at 37°C for the indicated period of time. To follow Toxoplasma multiplication EGFP expressing parasites were allowed to infect MDMs (MOI 1-3) in black 96-well plates. After 45 min cells were washed before adding RPMI with 10% FCS and Tats as indicated but without phenol red. EGFP fluorescence was quantified after 24h using a Tecan Spark 10 plate reader.

### Infection with Mtb

*M. tuberculosis* H37Rv-dsRed strain was grown in suspension in Middlebrook 7H9 medium (BD) supplemented with 10% albumin-dextrose-catalase (ADC, BD), 0.05% Tween-80 (Sigma-Aldrich) and 100 µg/mL Hygromycin B (InvivoGen) [75]. For infection, growing Mtb at exponential phase was centrifuged at 3000 g and resuspended in Phosphate-Buffered Saline (PBS, Gibco) (MgCl_2_ & CaCl_2_ free). Bacterial aggregates were dissociated by twenty passages through a 26G blunt needle. Bacterial suspension concentration was determined by measuring the optical density at 600 nm and then resuspended in PBS for in vivo infection. MDMs pretreated with Tat were infected with M. tuberculosis H37Rv-dsRed (MOI=0.3). After 18 h of infection at 37°C, the medium was changed and cells were fixed for analysis at 1, 3 and 5 days post-infection.

### Western blots

hMDMs were pretreated with 15 nM Tat for 5 h at 37°C. When indicated 20 nM bafilomycin A1 was then added for 1 h. Macrophages were then incubated for 20 min at 4°C in PBS supplemented with 5 mM EDTA. Cells were then scraped, centrifuged and stored as dried pellets at -80°C. Cells were resuspended in SDS/PAGE reducing sample buffer, briefly sonicated then heated 3 min at 96% before loading on 10 % tricine or tris/ glycine gels to quantify LC3 and perilipin2, respectively. Gels were blotted on PVDF or nitrocellulose membrane for labelling LC3 and perilipin2, respectively. Membranes were blocked in Tris-buffered saline containing 0.1% Tween and 5 % dried skimmed milk. This solution was used to dilute primary and secondary antibodies. LC3 blots were stained with HRP-secondary antibodies for imaging using a Biorad Chemidoc, while perilipin2 blots were labelled with IR dye 800 for Odyssey imaging.

### Imaging

hMDMs were pretreated with recombinant Tat for 5 h at 37°C. Lipid droplets were labeled with BODIPY 493/503 as described [62]. Intracellular labelling was quantified using Fiji. To label LC3B, cells grown on coverslips were washed then fixed in acetone/methanol at -20°C for 5 min and washed with ice-cold PBS. Anti-LC3B was then added in permeabilizing solution (PBS supplemented with 3 % FCS, 1 mg/ml FCS and 0.5% saponin) for 1 h at RT. After washes with permeabilizing solution, donkey anti-mouse-Cy3 was added for 1h. Cells were then washed, labeled with 500 nM DAPI in PBS, then mounted in Vectashield plus (Vector laboratories). For p62 staining, hMDMs were grown on coverslips, then infected with drRed-expressing H37Rv Mtb. After 1-2 days they were fixed with 3.7% paraformaldehyde, then permeabilized with Triton X-100 0.3% (Sigma-Aldrich) for 10 min and blocked with BSA (1% in PBS) for 30 min. Cells were incubated with anti-p62 antibodies for 1 hour, washed in PBS then incubated with AlexaFluor488 conjugated anti-mouse antibodies (Cell Signaling Technology) and DAPI (500 ng/mL, Sigma Aldrich) for 30 min. After several washes with PBS, coverslips were mounted on a glass slide using fluorescence mounting medium (Dako) and stored at 4°C.

Cells were examined using a Zeiss LSM 880 confocal microscope and a 63x objective. Representative median optical sections are shown. Image analysis was performed using Fiji. Vacuoles containing *T.gondii* and Mtb were identified using EGFP fluorescence and DAPI staining or dsRed, respectively. For quantification, labeling of these vacuoles with LC3B was examined for at least 30 cells.

To follow transferrin uptake by hMDMs, cells were pretreated with 15 nM Tat then labelled with 100 µM transferrin-FITC for 5 min in RPMI supplemented with 1% FCS, before cell fixation, confocal microscopy and quantification of intracellular and plasma membrane labeling for n>50 cells.

Autophagosomes in hMDMs were identified using anti-LC3 labeling. They were quantified on LSM880 confocal images either by counting the number of spots or by measuring the percentage of cell area occupied by these spots using the Fiji particle analysis. A threshold of 3 pixels was set for particle size. At least 30 cells were analyzed.

Autophagosomes in RAW macrophages were followed after transfection with mCherry-LC3B. When indicated, cells were co-tranfected (1/1) with Tat or dynamin mutants or treated with 30 µM Pitstop2 or Dyngo4a for 90 min. Cells were then labelled for 30 min with 100 µM transferrin-Cy5, fixed, labelled with DAPI, mounted then examined using a Spinning disk Nikon Ti Inscoper CSU-X1 with a 100 x 1.45 NA objective. LC3 dots representing autophagosomes (size> 3 pixels) were then counted and transferrin internalization efficiency was measured as described above.

Live imaging of RAW macrophages transfected with mRFP-GFP-LC3 was performed at 37°C in RPMI/FCS without phenol red under 5% CO2 using a Spinning disk Nikon Ti Inscoper CSU-X1 microscope.

Colocalization between green and red spots was quantified using the Mander’s coefficient calculated using the JACoP plugin of Fiji.

For TIRF imaging RAW macrophages were grown on 4-well glass-bottom Ibidi µ-slides (#80427) and transfected with AP-2-σ2-EGFP or the indicated EGFP-tagged protein. When indicated, cells were treated with 15 nM recombinant Tat for 5 h. They were then fixed with paraformaldehyde, washed and kept in PBS for imaging using a Nikon inverted microscope fitted with a 100x NA1.49 objective and a iLAS2 Roper TIRF illumination module. The percentage of cell area occupied by EGFP spots was quantified using the Fiji particle analysis.

### Zebrafish experiments

#### Zebrafish husbandry and ethic statements

All zebrafish (*Danio rerio*) procedures were performed by authorized staff and conducted by following the 3Rs (-Replacement, Reduction and Refinement-) principles in compliance with the European Union guidelines for handling of laboratory animals. Local standards set were approved by the Direction Sanitaire et Vétérinaire de l’Hérault and the Comité d’Ethique pour l’Expérimentation Animale de la Région Languedoc Roussillon and the French Ministry of Agriculture (authorization number: APAFIS #36309-2022040114222432). Breeding and maintenance of adult zebrafish were performed at the ZEFIX (Lphi, UMR5294 CNRS) Home Office-approved aquaria, according to the local animal welfare standards (license number CEEA-LR-B34-172-37). Experiments were performed using the wild type AB line. The number of animals used for each procedure was guided by pilot experiments. For zebrafish anesthesia procedures, larvae are immersed in a 0.168 mg/mL Tricaine (Sigma-Aldrich) solution in fish water. When required, larvae were cryo-anesthetized by incubation on ice for 10 min then euthanized using an overdose of Tricaine (0.5 mg/mL).

#### Bacterial strain and Growth conditions / Mycobacterium marinum

(M strain) carrying pTEC27 (Addgene, plasmid 30182) that express a red fluorescent protein (tdTomato) was used for this study. *M. marinum* expressing tdTomato were grown at 28.5 °C under hygromycin B selection (50 µg/ml, Invivogen) in Middlebrook 7H9 broth medium supplemented with albumin, dextrose, catalase (ADC, Millipore-Sigma). To prepare *M. marinum* inoculates, mid-log-phase cultures of *M. marinum* were pelleted, washed and resuspended in PBS. Mycobacterial suspensions were then homogenized through a 26-gauge needle, harvested by centrifugation and resuspended in a volume of PBS appropriate to the bacterial dose required then kept on ice until cellular or zebrafish infection challenges.

In vivo *mycobacterial infection, treatment and analysis.* Microinjections of Tat (6 nL) were carried out into the caudal vein in 30 hpf embryos previously dechorionated and tricaine-anesthetized. Treated-embryos were then directly intravenously infected with 2 nl of mycobacterial suspensions (approximately 150 mycobacteria). The inoculum size was checked by injection of 2 nL in sterile PBS and plated on 7H10 ADC agar (Millipore-Sigma). Groups of infected/treated embryos were then transferred into plates and incubated at 28.5 °C. Tat administration in *M. marinum*-infected animals was repeated at 1 and 2 days post infection. Survival post *M. marinum* infection was assessed daily by counting dead embryos (no heartbeat) up to 10 days. In addition, to determine effects of Tat on infection outcomes, larvae were collected at 2, 3 and 4 days post infection and imaged for both granuloma quantification (defined at least 10 infected cells) and bacterial burdens analysis as Fluorescent Pixel Count (FPC) by fluorescence microscopy (MVX10 Olympus microscope, MVPLAPO 1X objective; XC50 camera) [76].

## Acknowledgements

We thank Simon Lachambre, Gustavo Stadthagen-Gomez and Hélène Botella for help during pioneer experiments, Laurent Kremer, Wassim Daher, Alenka Copic, Maria Morel-Carretero and Laura Picas for discussions, and all the people that provide valuable reagents for this study. We acknowledge the ZEFIX-LPHI (ZEbraFIsh and Xenopus platform, Lphi, University of Montpellier) aquarium teams for zebrafish maintenance and care and the team of the Montpellier Ressources Imagerie facility for help with imaging. This work was funded by Sidaction (grant # 23-1-AEQ-13605-1) to BB and CV, and partly by a grant from the Agence Nationale de Recherche sur le Sida et les hépatites virales (ANRS) to ON and CV. SB acknowledges funding from the Agence Nationale de la Recherche (grant ANR-24-CE15-3620).

## Legends to Figures

**Figure S1.**
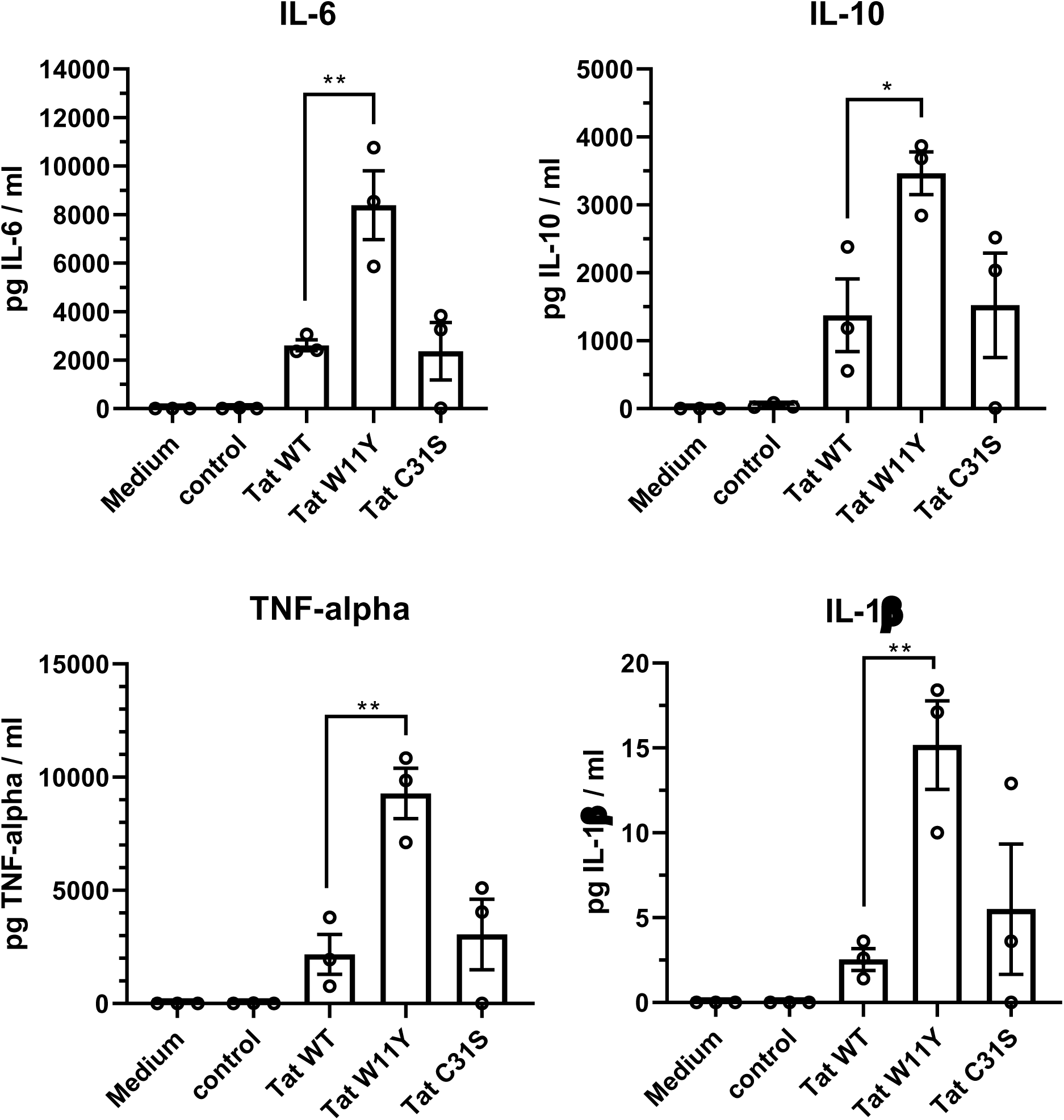
Tat WT, Tat-W11Y and Tat-C31S induce cytokine secretion. hMDMs were treated with 15 nM Tat for 5 h before harvesting supernatant for cytokine assays. Data are mean ± SEM (n=3) of cells from three different donors. One-way ANOVA compared to Tat WT (*, p<0.05; **, p<0.01).

**Figure S2.**
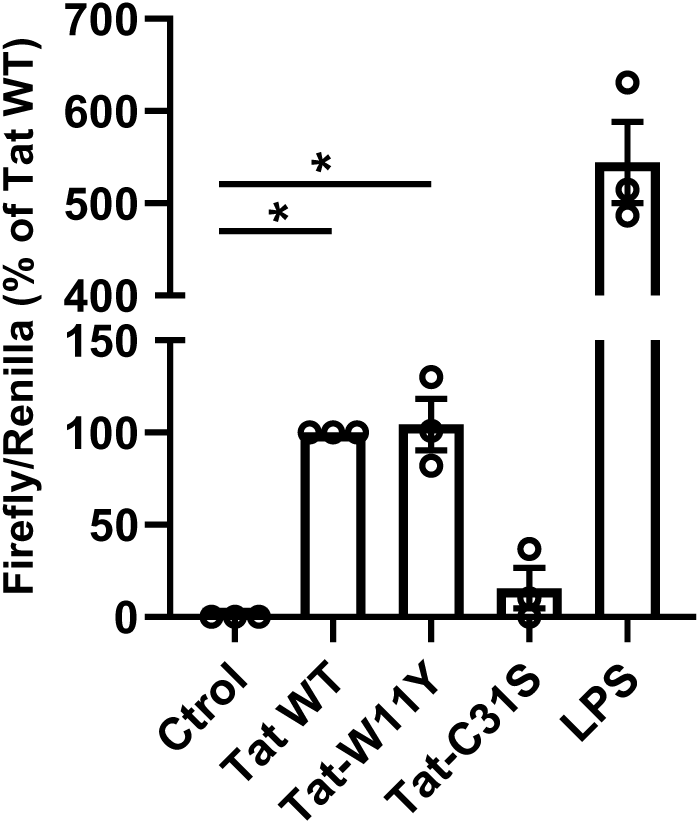
Tat WT and Tat-W11Y but not Tat-C31S activate NF-κB. RAW. 264.7 cells were transfected with a vector containing the Firefly gene behind 3 NF-κB sites and another expressing the Renilla luciferase. Cells were then treated for 5 h with 15 nM Tat (WT, W11Y or C31S as indicated) or 0.5 µg/ ml of E coli LPS as positive control, before harvesting cells for luciferase assays, Results (mean ± SEM, n=3) are expressed as Firefly / Renilla activity ratio. One-way ANOVA compared to control (*, p<0.05).

**Figure S3.**
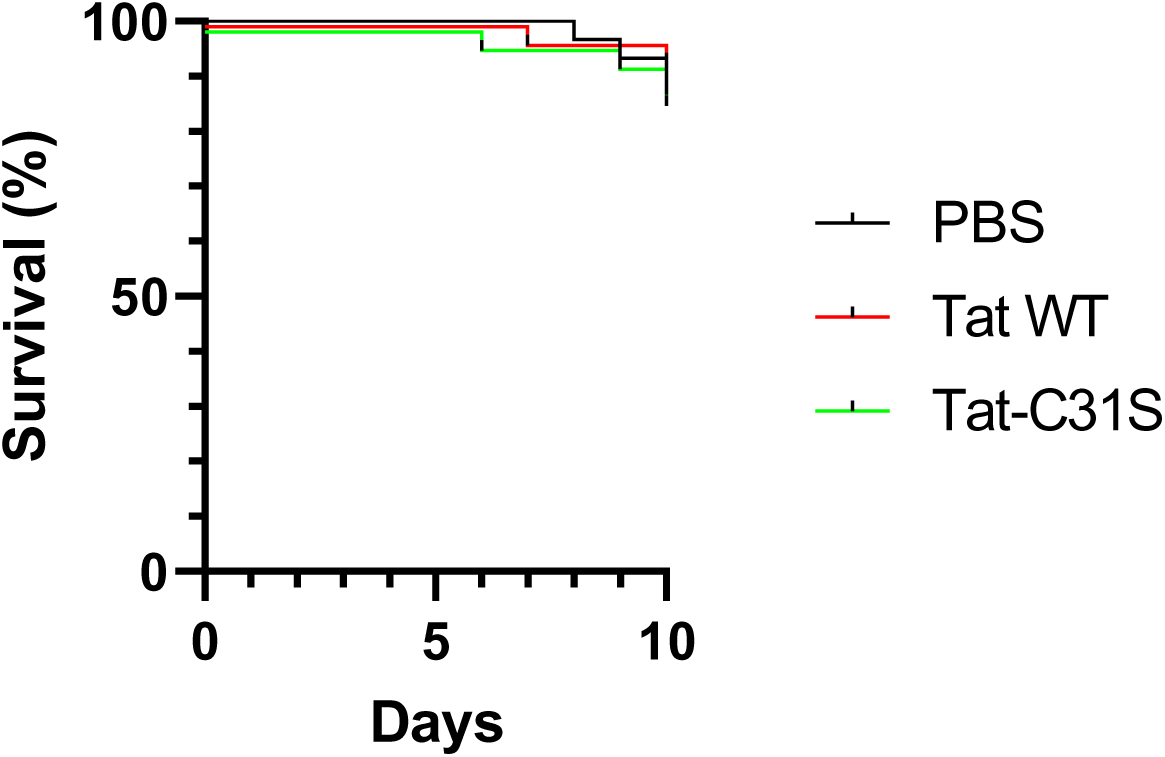
Tat injections do not affect the survival of zebrafish embryos. Zebrafish embryos (n=20-25 for each group) at 24 hpf were injected with Tat (∼100 nM final concentration) WT or C31S, as indicated, Injections were repeated at day 2,3,4 and 5.

**Figure S4.**
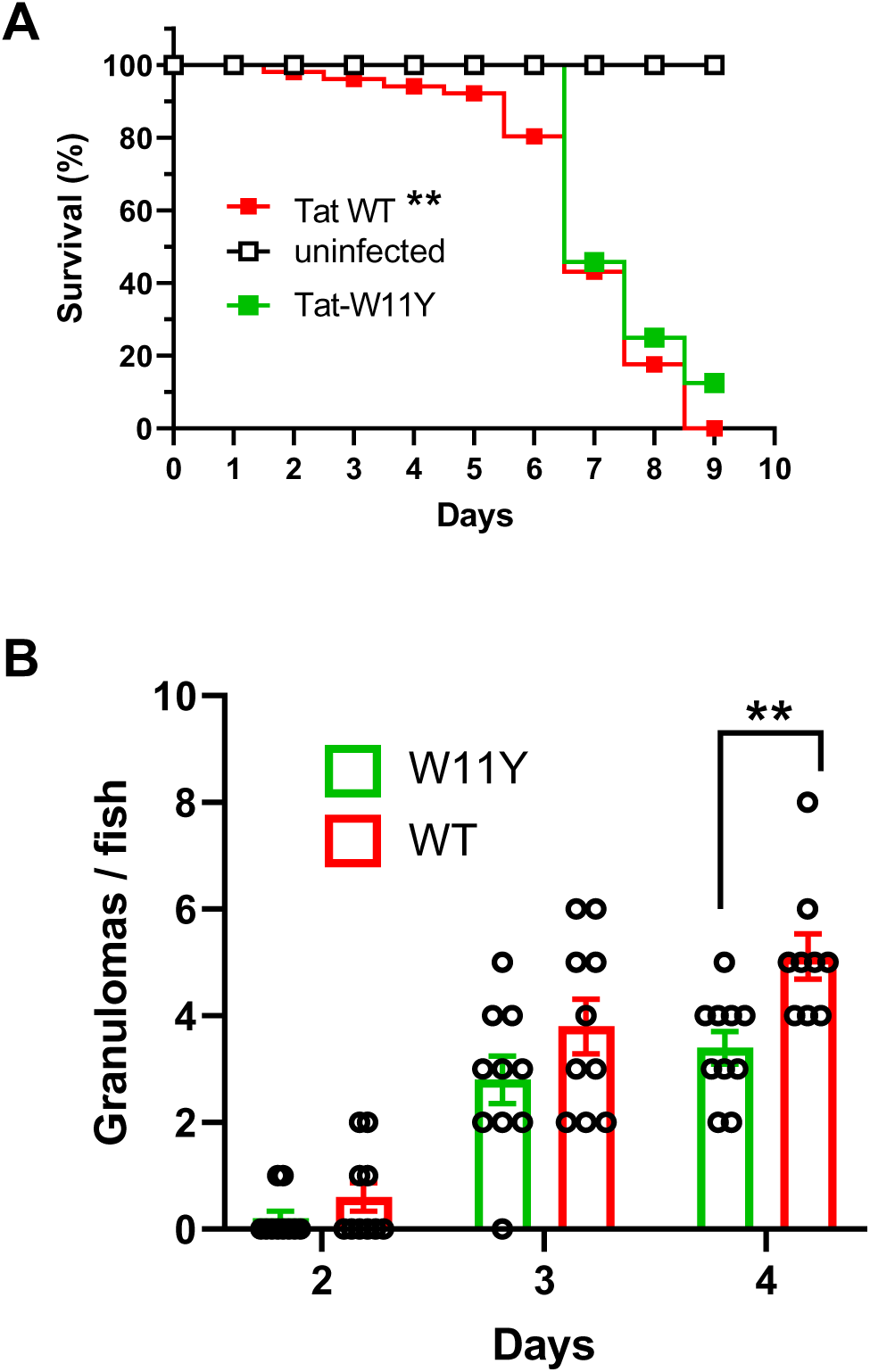
Effect of Tat W11Y on the multiplication of Mycobacterium marinum in Zebrafishes. A, Zebrafish embryos (n=20-25 for each group) at 24 hpf were injected with Tat (∼100 nM final concentration) then infected with *Mycobacterium marinum.* Tat injection (WT or W11Y) was repeated at dpi 2,3,4 and 5. Survival curves of the Tat groups (WT/ W11Y) were compared using Log-rank Test (**, p<0.01). B, zebrafishes were infected with tdTomato-M. marinum, Granulomas were counted after the indicated number of dpi. Two ways ANOVA (**, p<0.01).

**Figure S5.**
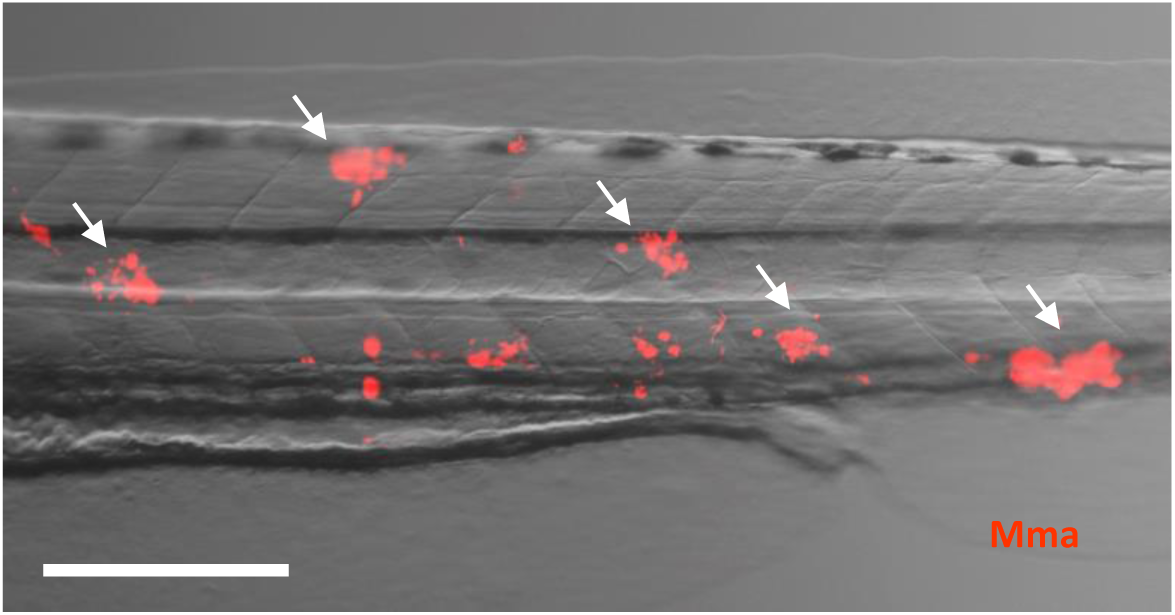
Examples of granulomas in Zebrafish embryos. Zebrafish embryos at 24 hpf were infected with tdTomato *Mycobacterium marinum* (Mma), and imaged 3 dpi. Arrows point to granulomas. Bar, 200 µm.

**Figure S6.**
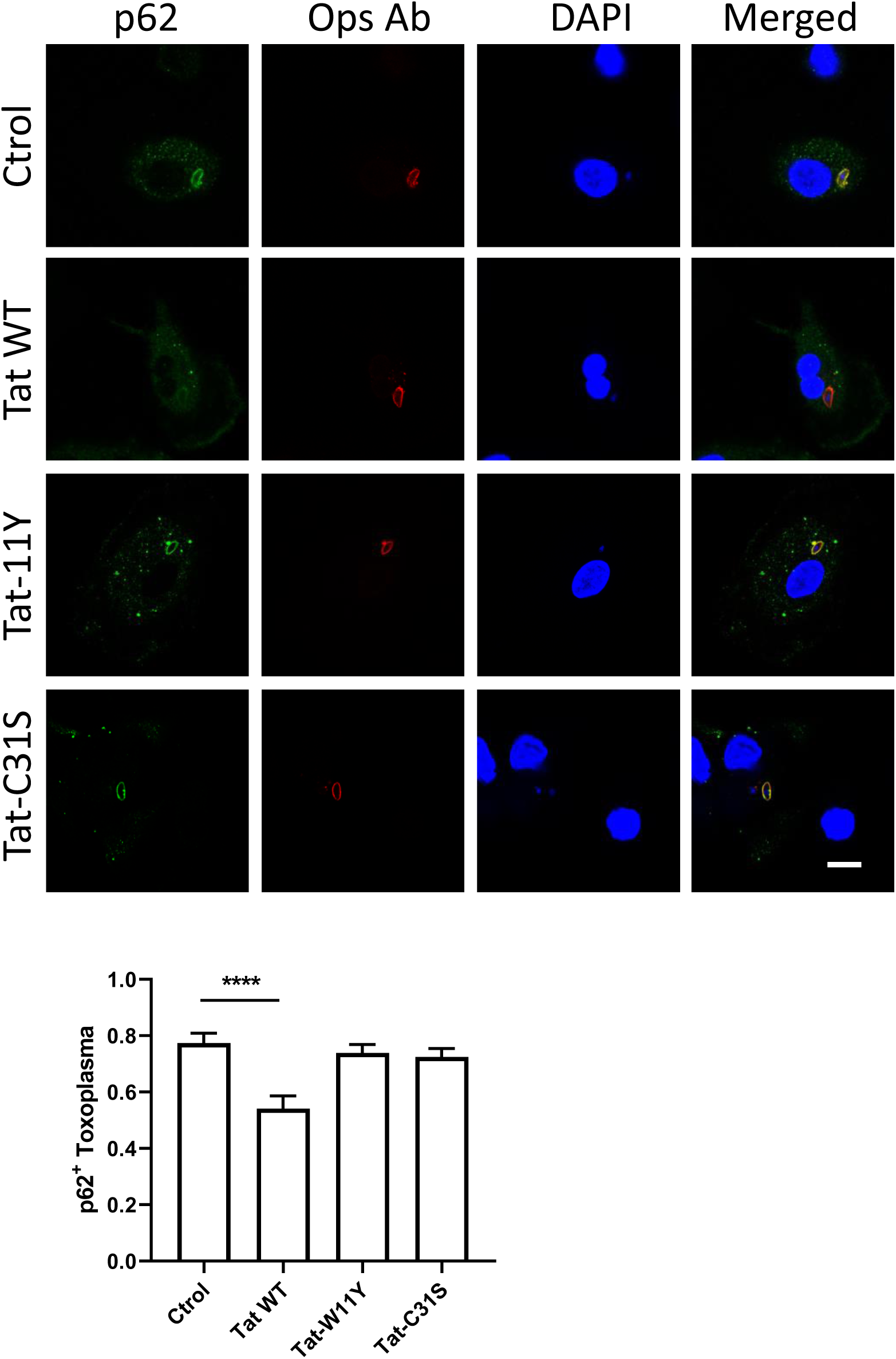
Tat inhibits the recruitment of p62 on vacuoles containing *T. gondii*. hMDMs were pretreated with 15 nM Tat for 5 h, then infected with opsonized T. gondii (MOI=10) for 30 min before staining for p62 and opsonizing antibody, and DAPI staining, Representative confocal sections are shown. Bar, 10 µm. The graph shows the quantification of the fraction of P62^+^ toxoplasma on 250-500 parasites. One Way ANOVA, ***, p<0.0001

**Figure S7.**
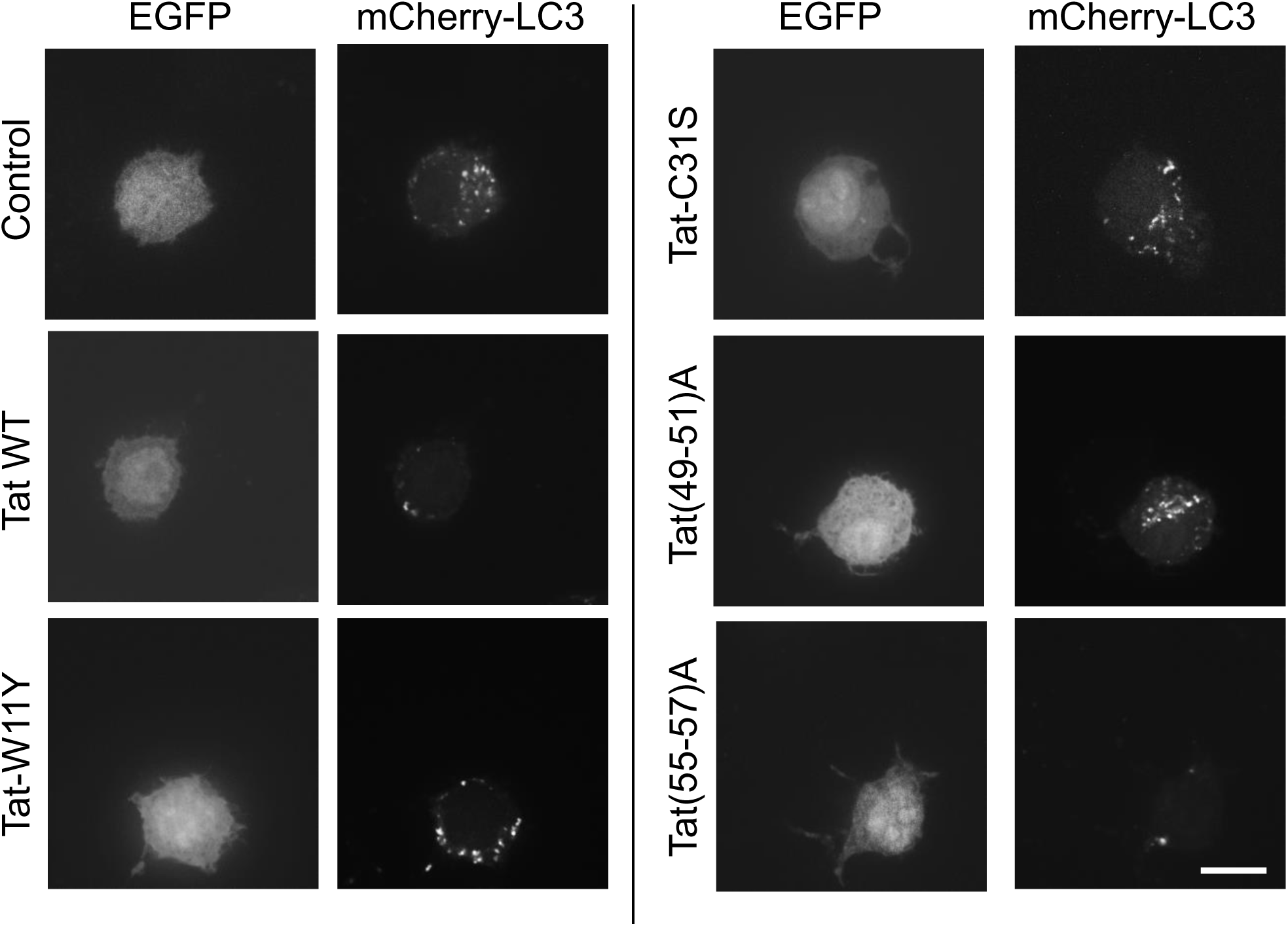
Effect of Tat mutants on autophagosome numbers. RAW macrophages were transfected with mCherry-LC3 and the indicated version of Tat. The Tat expression vector is bicistronic and also expresses EGFP. Cells were fixed after 18h and imaged using a spinning disk confocal microscope and a x100 NA 1.45 objective. Bar, 10 µm. Less autophagosomes are present in cells transfected with Tat WT and Tat(55-57)A.

**Figure S8.**
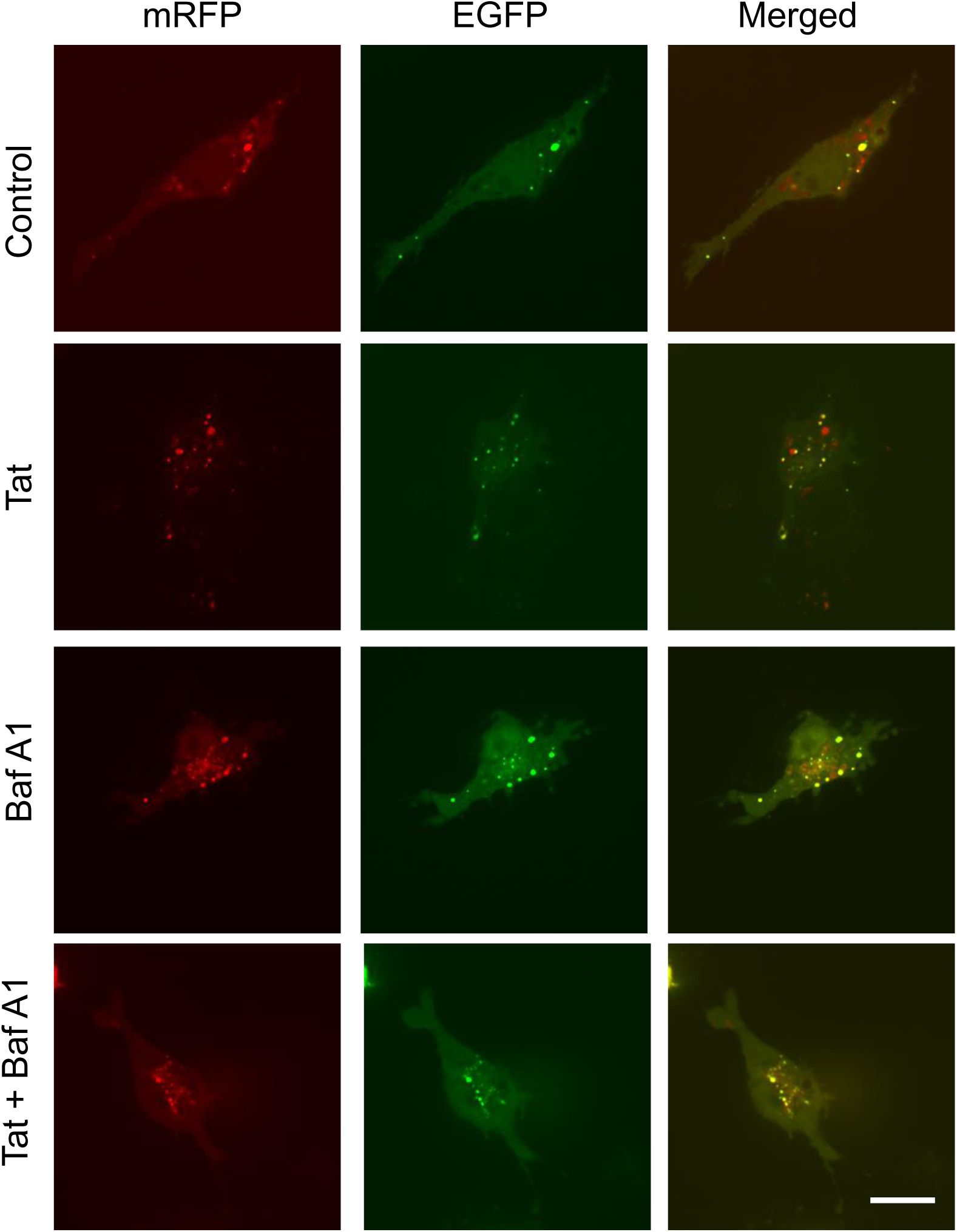
Tat does not affect autophagic flux, as monitored using mRFP-EGFP-LC3. RAW macrophages were transfected with mRFP-EGFP-LC3 then treated with 15 nM Tat for 4 h. When indicated, 20 nM Bafilomycin A1 was added for 1 h and cells were imaged at 37°C using a spinning-disk confocal microscope. Bar, 10 µm. Green structures accumulate when Baf is present.

**Figure S9.**
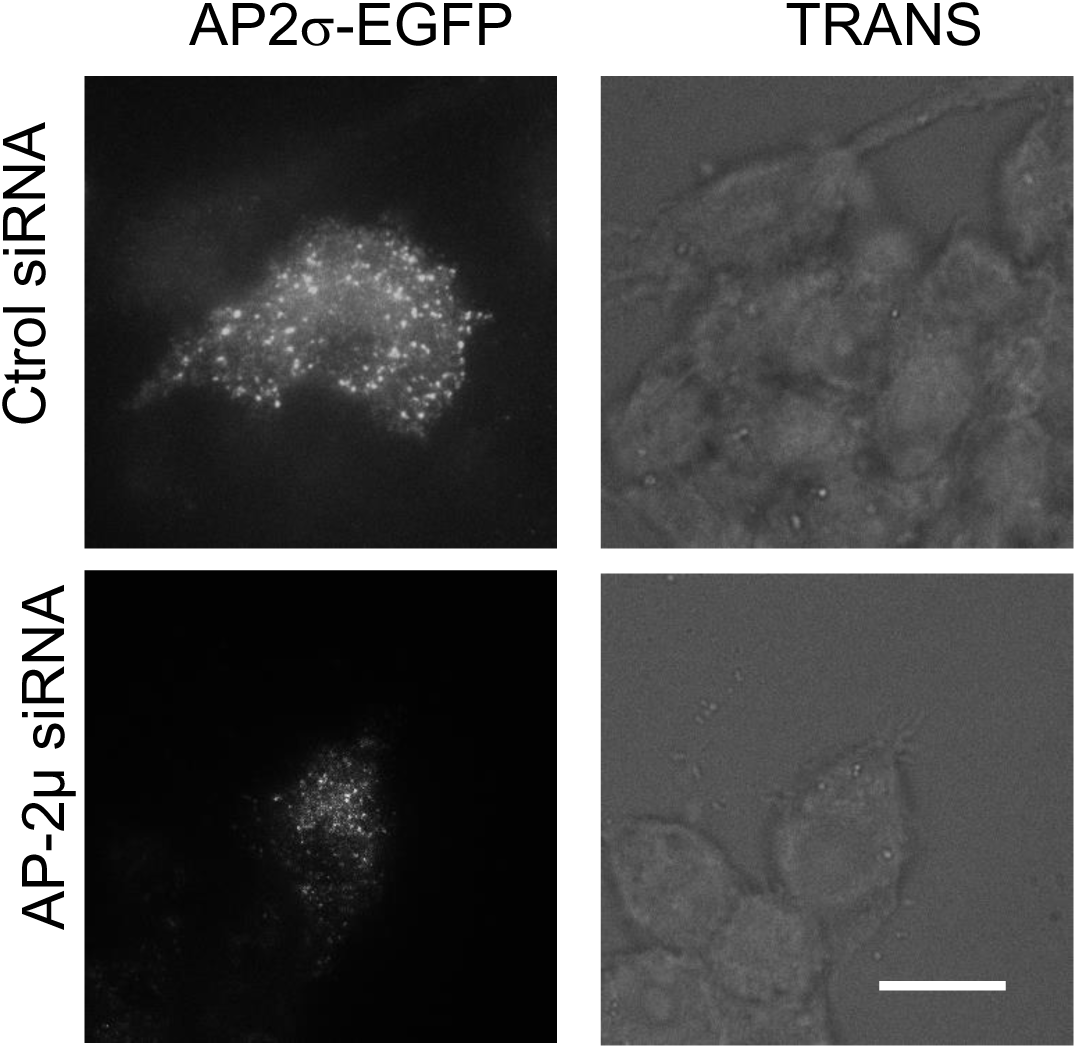
AP-2µ siRNA inhibits AP-2α2-EGFP recruitment at the plasma membrane. RAW cells were cotransfected with AP-2α2-EGFP and the indicated siRNA before fixation and imaging by TIRF microscopy using a x100 NA and a 488 nm laser for EGFP fluorescence or a LED for transmitted light (TRANS). Bar, 10 µm.

**Figure S10.**
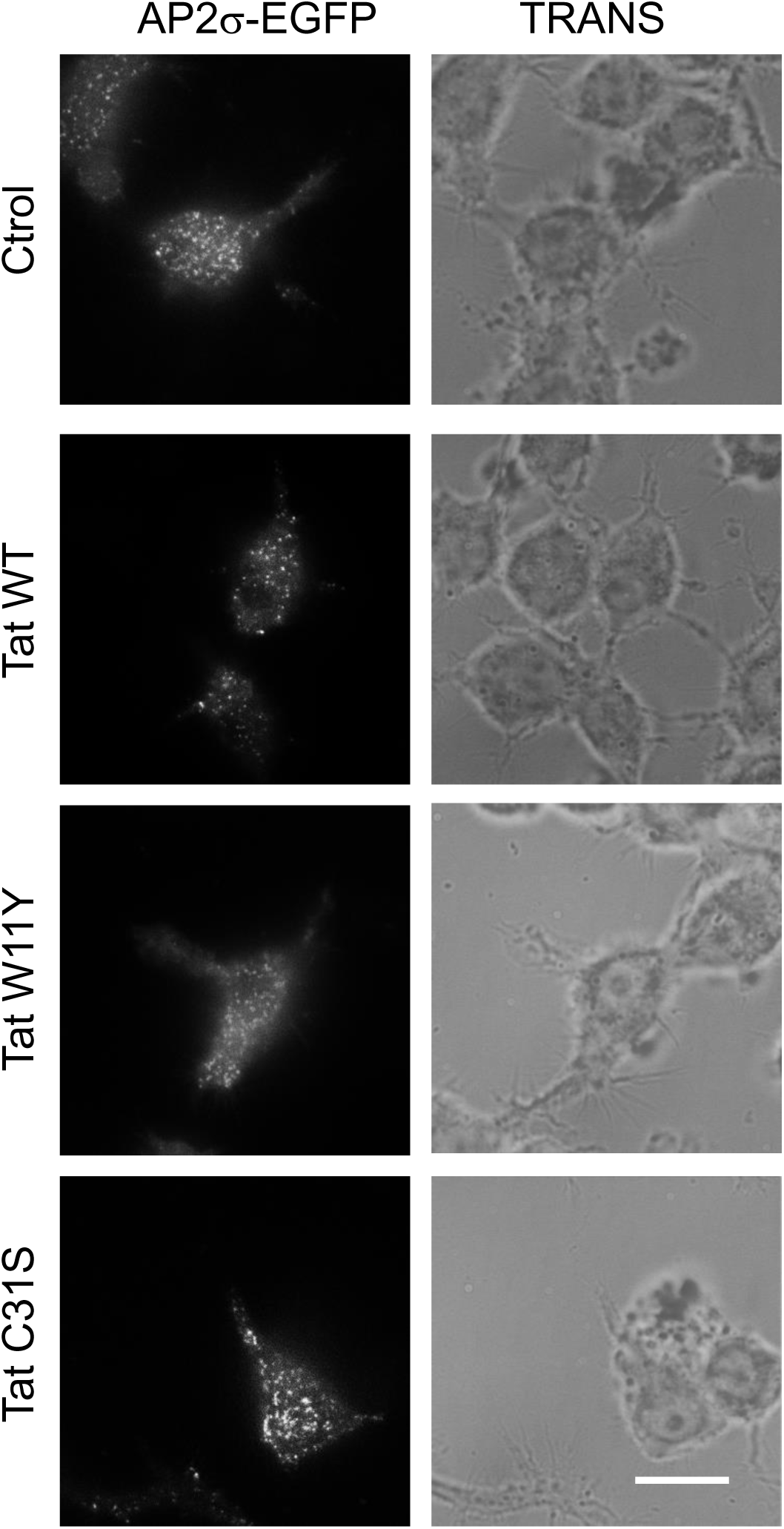
Tat palmitoylation is required for Tat to inhibit AP-2 recruitment. RAW macrophages were transfected with AP-2α2-EGFP, treated with 15 nM of the indicated Tat mutant for 5h before fixation and imaging by TIRF microscopy using a 488 nm laser for EGFP fluorescence or a LED for transmitted light (TRANS). Bar, 10 µm.

**Figure S11.**
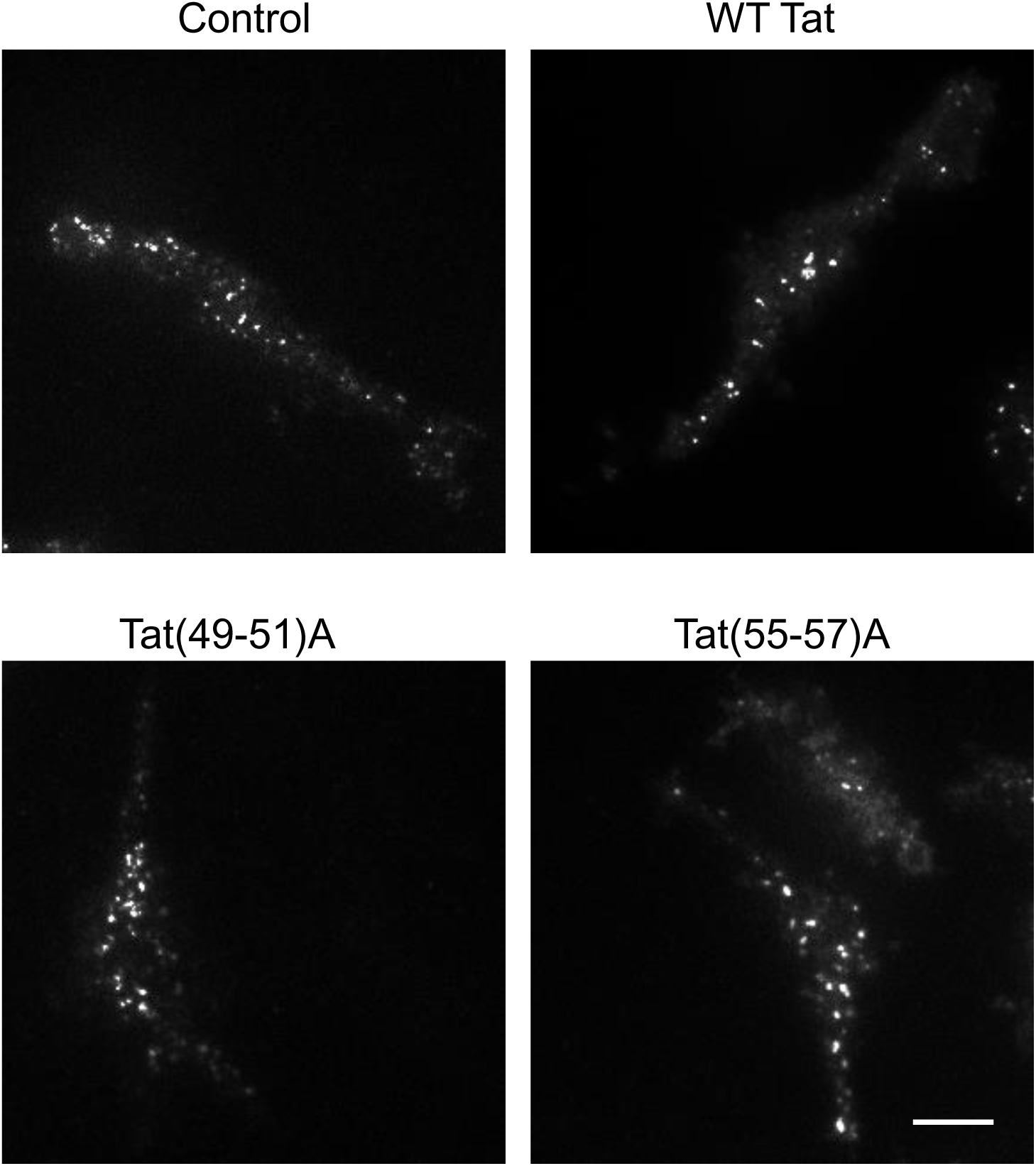
Tat Binding to PI(4,5)P2 is required for Tat to inhibit AP-2 recruitment. RAW 264,7 macrophages were cotransfected with AP-2s2-EGFP and the indicated Tat mutant before fixation and imaging by TIRF microscopy. Bar, 5 µm.

**Figure S12.**
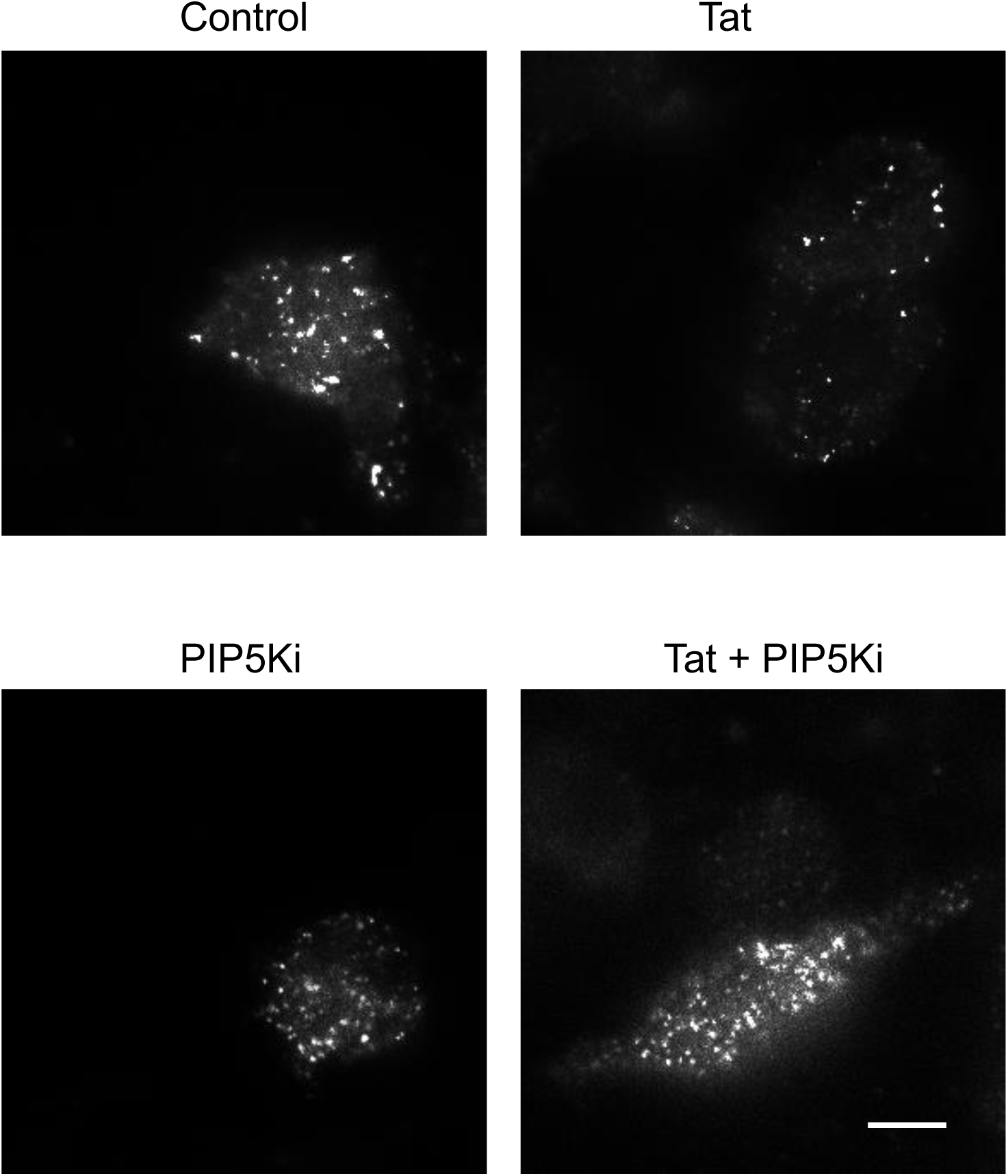
Increasing PI(4,5)P2 level prevents Tat from inhibiting AP-2 recruitment. RAW macrophages were cotransfected with AP-2α2-EGFP, Tat and the mouse PIP-5 kinase as indicated before fixation and imaging by TIRF microscopy. Bar, 5 µm.

**Figure S13.**
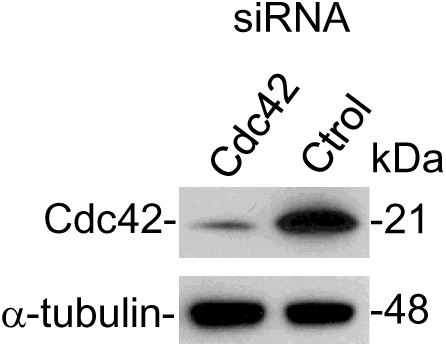
An siRNA against Cdc42 inhibits its expresssion. RAW 264.7 macrophages were transfected using RNAiMAX with a control siRNA, or an siRNA against Cdc42. After 24h, cells were lysed before western blotting against Cdc42 or α-tubulin as a loading control.

**Figure S14.**
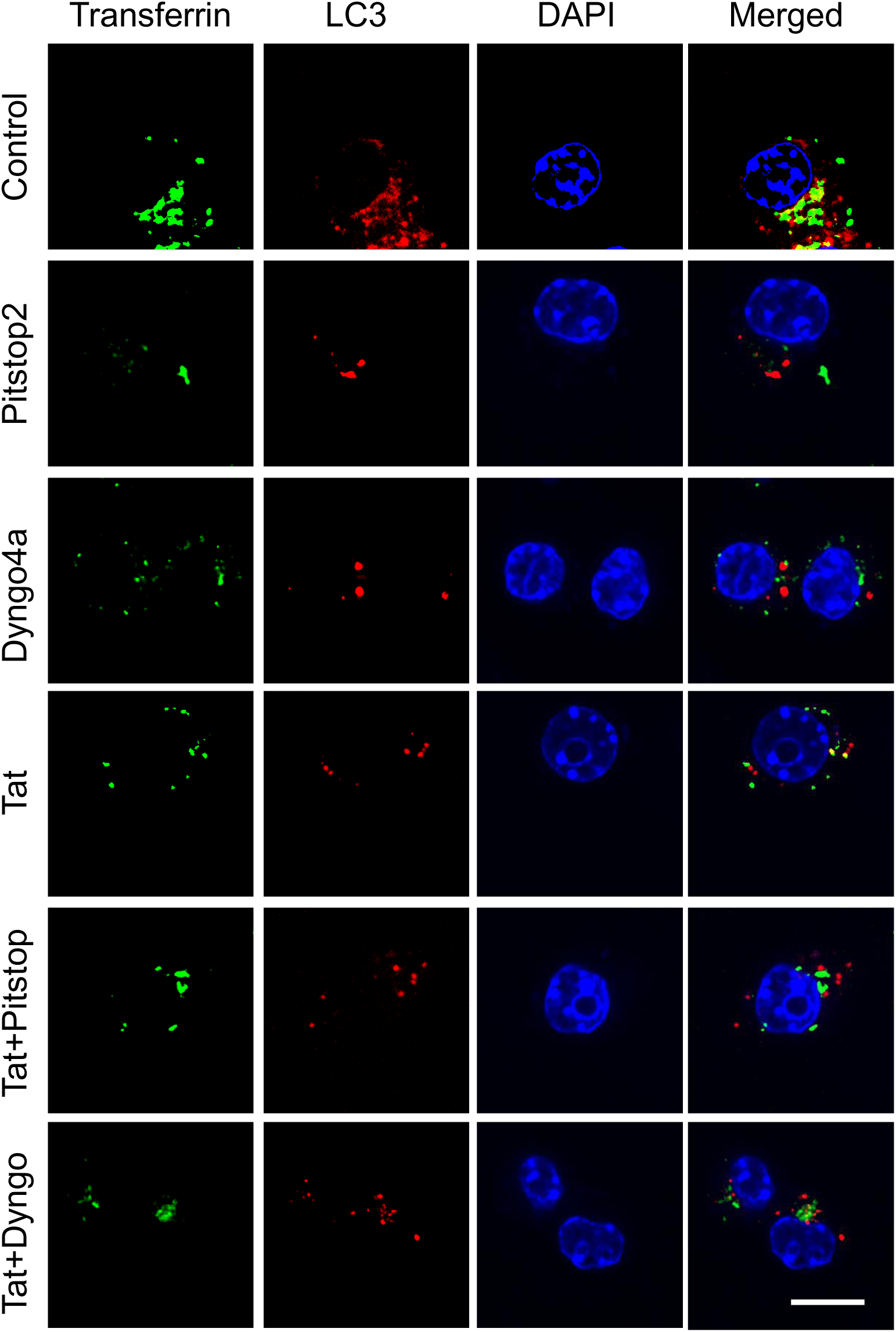
CME inhibitors and Tat do not show additive inhibitory effects on CME and autophagy. RAW macrophages were transfected with mCherry-LC3 and Tat as indicated, treated with 30 µM Pistop2 or Dyngo4a for 90 min before labeling cells with Cy5-transferrin for 30 min, fixation, and confocal microscopy. Bar, 10 µm.

**Figure S15.**
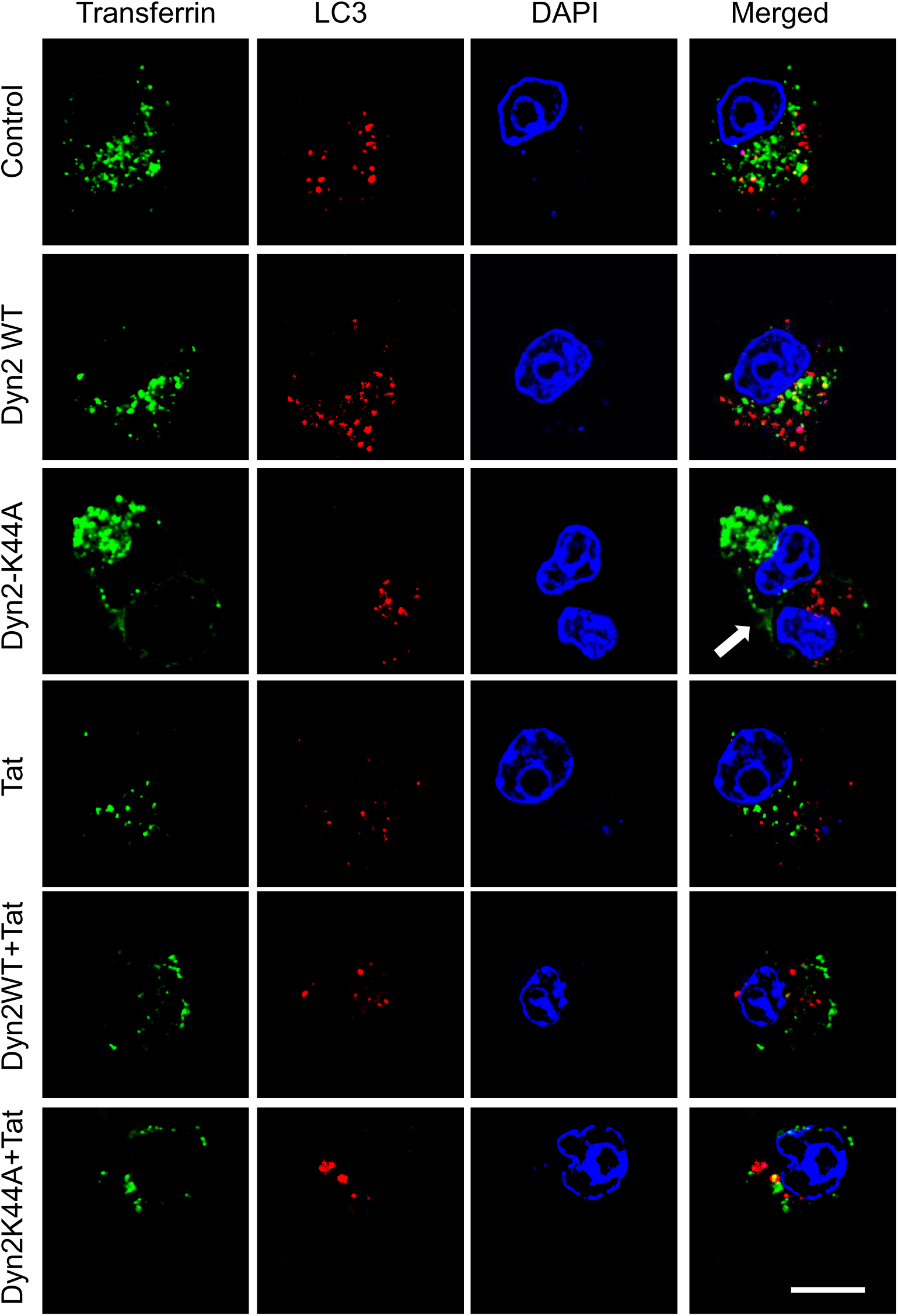
Dominant negative dynamin2 and Tat do not show additive inhibitory effects on CME and autophagy. RAW cells were transfected with mCherry-LC3, Dynamin2 (WT or K44A) and Tat as indicated before labeling cells with Cy5-transferrin for 30 min, fixation, and confocal microscopy. Arrow points at transfected cell when needed. Bar, 10 µm.

**Figure S16.**
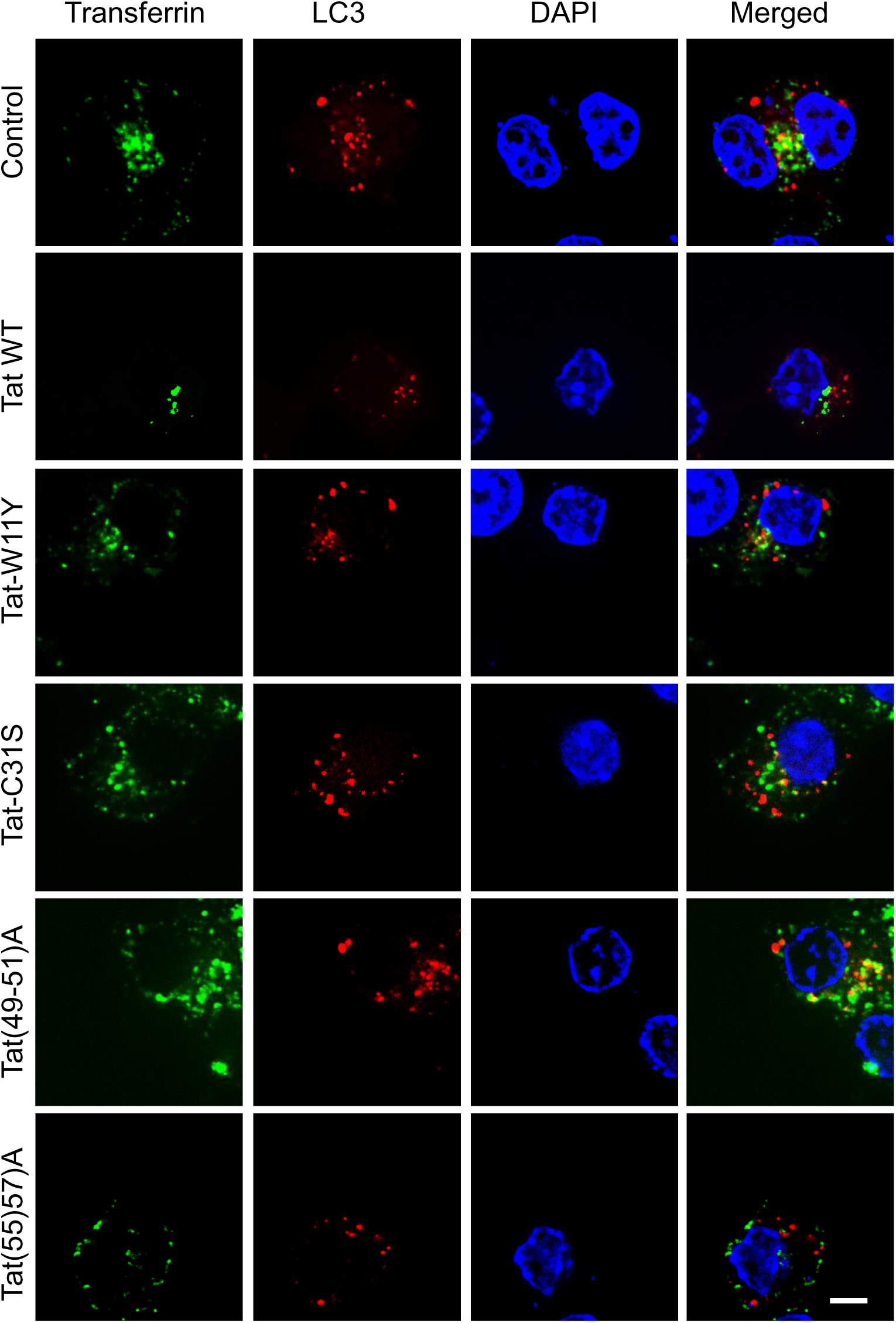
Tat binding to PI(4,5)P2 is required for Tat to inhibit CME and autophagy. RAW macrophages were transfected with mCherry-LC3 and the indicated Tat mutant before labeling cells with Cy5-transferrin for 30 min, fixation, and confocal microscopy. Bar, 5 µm.

**Figure S17.**
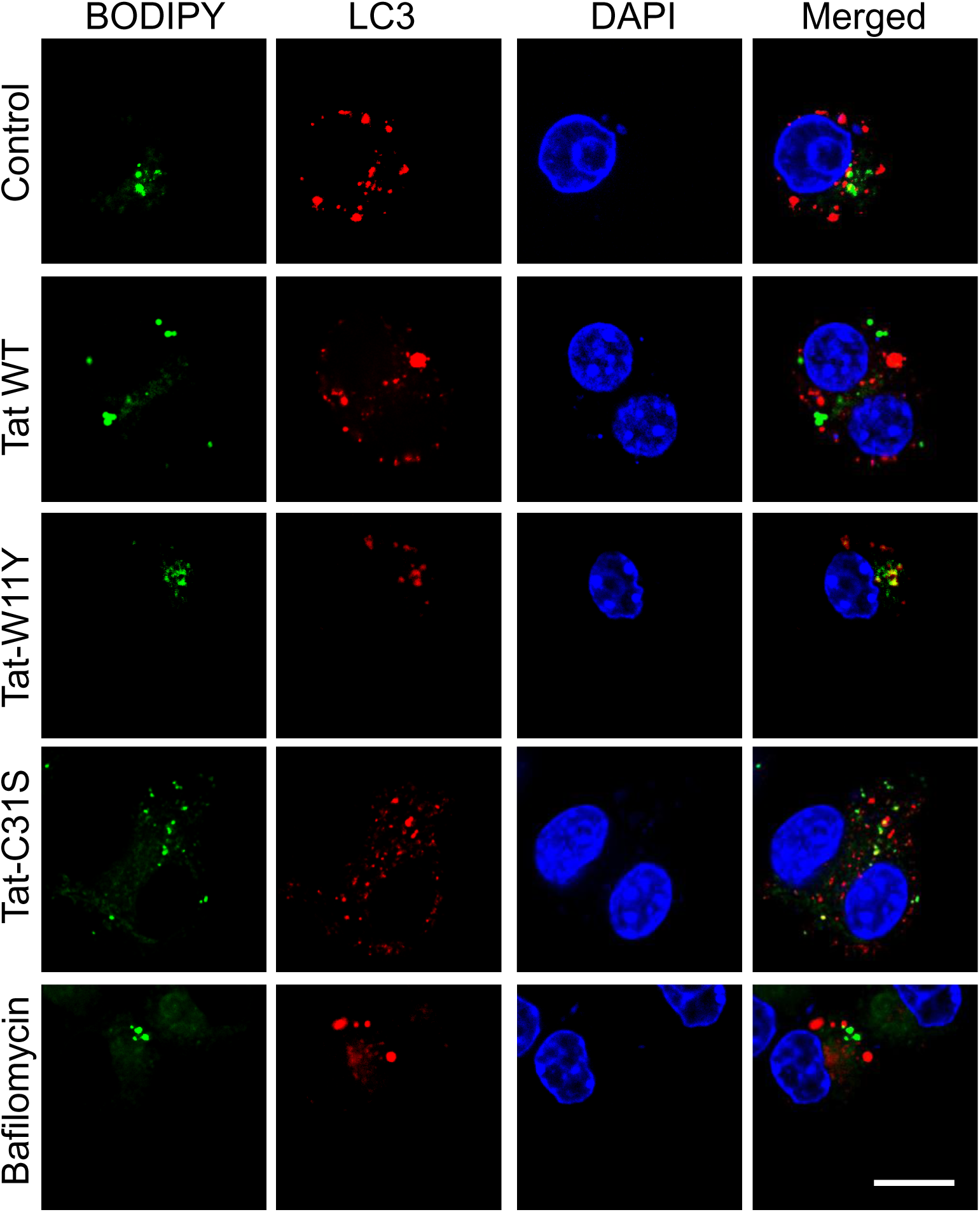
Tat decreases the localization of lipid droplets in LC3-positive structures. RAW 264.7 cells were transfected with mCherry-LC3, treated with 15 nM of the indicated Tat mutant for 5 h or 100 nM bafilomycin A1 for 2 h before fixation, staining with BODIPY 493/503 and DAPI, and imaging by confocal microscopy. Bar, 10 µm.

